# Heterologously secreted MbxA from *Moraxella bovis* induces a membrane blebbing response of the human host cell

**DOI:** 10.1101/2022.05.02.490299

**Authors:** Isabelle N. Erenburg, Sebastian Hänsch, Feby M. Chacko, Anna Hamacher, Sebastian Wintgens, Fabian Stuhldreier, Gereon Poschmann, Olivia Spitz, Kai Stühler, Sebastian Wesselborg, Johannes Hegemann, Sander H. J. Smits, Stefanie Weidtkamp-Peters, Lutz Schmitt

## Abstract

Many proteins of the Repeats in Toxins (RTX) protein family are toxins of Gram-negative pathogens including hemolysin A (HlyA) of uropathogenic *E. coli*. RTX proteins are secreted via Type I secretion systems (T1SS) and adopt their native conformation in the Ca^2+^-rich extracellular environment. Here we employed the *E. coli* HlyA T1SS as a heterologous surrogate system for the RTX toxin MbxA from the bovine pathogen *Moraxella bovis*. In *E. coli* the HlyA system successfully activates the heterologous MbxA substrate by acylation and secretes the precursor proMbxA and active MbxA allowing purification of both species in quantities sufficient for a variety of investigations. The activating *E. coli* acyltransferase HlyC recognizes the acylation sites in MbxA, but unexpectedly in a different acylation pattern as for its endogenous substrate HlyA. HlyC-activated MbxA shows host species-independent activity including a so-far unknown toxicity against human lymphocytes and epithelial cells. Using live-cell imaging, we show an immediate MbxA-mediated permeabilization and a rapidly developing blebbing of the plasma membrane in epithelial cells, which is associated with immediate cell death.

## Introduction

In light of a growing frequency of antibiotic resistance observed in human pathogenic bacteria, characterization of virulence factors, their posttranslational modifications and their secretion machineries are key factors for the development of novel anti-virulence strategies against bacterial infections. Targeting for example bacterial toxins is a promising approach for future therapeutics that need to be tailored to specific pathogens and pathogenicity factors (1, 2).

Proteins of the Repeats in Toxins (RTX) family are secreted to the extracellular space by many Gram-negative bacteria employing Type 1 Secretion Systems (T1SS). The growing number of sequenced bacterial genomes revealed novel members of this RTX family (3). Many RTX proteins are virulence factors of important pathogens such as the toxins CyaA from *Bordetella pertussis* or hemolysin A (HlyA) from uropathogenic *E. coli* (UPEC). RTX proteins are characterized by glycine reach nonapeptide repeats with the consensus sequence GGxGxDxU (x – any amino acid, U - large hydrophobic amino acid,) the so-called GG-repeats (3, 4). Repetitions of this repeat close to the C-terminus form the RTX domain (5), while the variability of the N-terminus between different RTX proteins allows diverse functions (for a review see (3)). One of the best-characterized T1SS is the HlyA secretion system from *E. coli*, which consists of the ABC transporter hemolysin B (HlyB), the membrane fusion protein hemolysin D (HlyD) and the outer membrane protein TolC (6–8). Here, the C-terminal, non-cleavable secretion sequence of 50-60 amino acids of HlyA (Fig. 1) conveys specific interaction with the inner membrane components HlyB and HlyD of this T1SS (9–12). Prior to and during transport, which occurs in a one-step mechanism across both membranes of the Gram-negative bacterium (9, 11, 12), HlyA remains unfolded and only achieves its native conformation upon secretion into the extracellular fluid, where binding of Ca^2+^ ions to the RTX domain induces folding of the protein (5, 13–15).

**Fig. 1:**
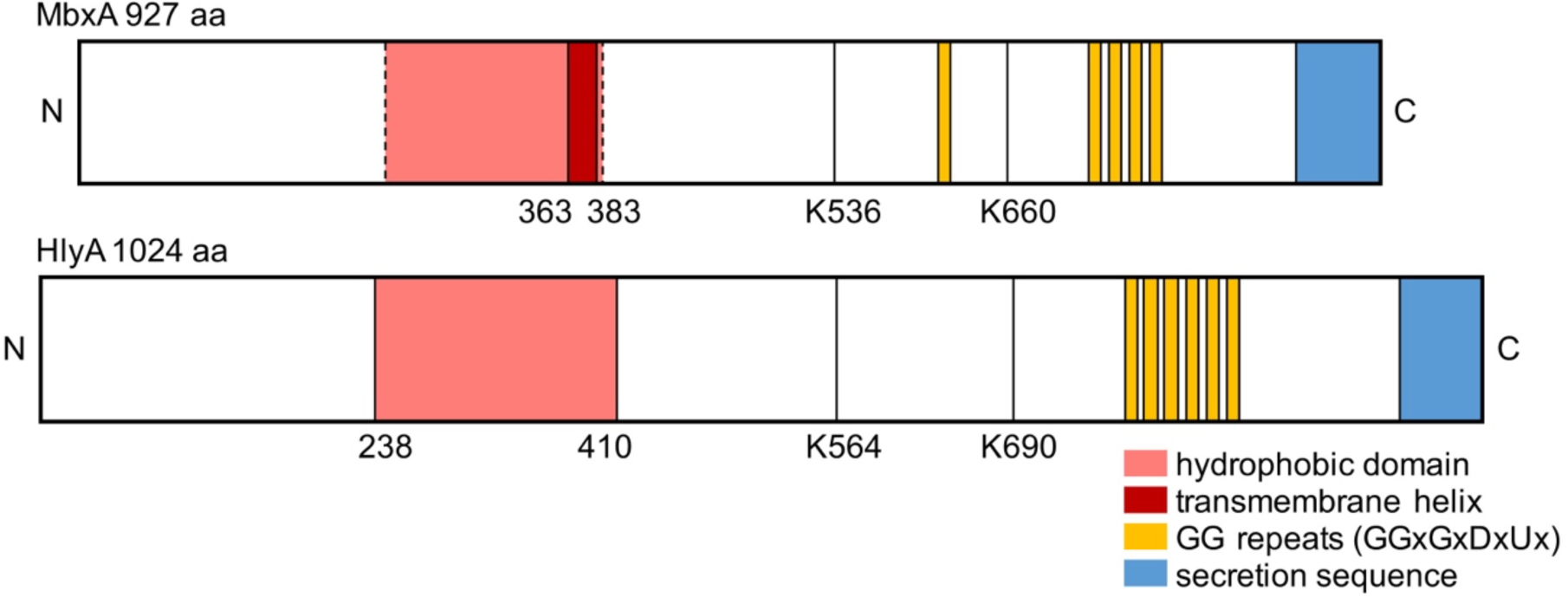
Schematic view of the primary structure of MbxA and HlyA. Five conserved GG repeats with the consensus sequence GGxGxDxUx are present in MbxA, while HlyA carries six conserved GG repeats (yellow boxes). They together form the Ca^2+^ binding RTX domain. As a RTX protein, HlyA characteristically possesses a C-terminal secretion signal of approximately 60 amino acids (10–12), which is shown in blue. For MbxA the length of the secretion sequence is unknown and drawn not to scale. In the N-terminal part of HlyA, a hydrophobic domain is involved in pore formation (53) and marked in red. Depending on the prediction algorithm (see Materials and Methods) for MbxA either (i) a single membrane spanning helix is predicted from residue 363 to 383, (ii) a set of transmembrane helices forming an ambiguous hydrophobic domain from residue 210 to 383, or (iii) even from residue 120 to 383 (dashed lines) are predicted (52). The position of the acylated lysine residues K564 and K690 of HlyA and the homologous residues predicted on sequence comparisons to become acylated in MbxA, K536 and K660, are indicated and served as anker point for the scheme.

HlyA belongs to the classic pore-forming RTX toxins and is active against a variety of cells including erythrocytes, leukocytes, epithelial and endothelial cells of different species (16–18). For the cytotoxic activity, but not for secretion, a posttranslational fatty acid acylation of two internal lysine residues (Lys 564 and Lys 690), catalyzed by the intracellular acyltransferase HlyC, is required prior to secretion (Fig. 1) (19, 20). Such an acylase is required for other RTX toxins such as CyaA from *Bordetella pertussis* or LktA from *Mannheimia haemolytica* as well and the gene encoding for the transferase is present in the operon coding for the T1SS. The structure of the acylase from *Actinobacillus pleuropneumoniae* (21) might serve as a blue print and provides first insights into how the acyltransferase transfers two fatty acids to the RTX toxin, a process for which acyl carrier protein (ACP) is required. For HlyA, the fatty acids transferred are predominantly C_14_ and to a minor extent C_15_ and C_17_ (22). Most importantly, both lysine residues can be acylated independent of each other *in vivo*, but the acylation of both lysine residues is strictly required for toxic activity of HlyA (20). The latter observation has been recently challenged (23).

Despite being studied for decades, the mechanism of pore-formation by RTX toxins remains elusive. Fatty acid acylation is required for cell lysis, but the architecture of the pore or the exact number of HlyA protomers forming the pore is unknown. Currently, two models are discussed. First, a pure proteinaceous pore, in which the pore lumen is formed exclusively by protein segments, and second a proteo-lipidic or toroidal pore, in which the pore lumen is formed by both, lipid and protein segments (24). The latter one is supported by recent data obtained for CyaA (25), while the first one was initially described by electrophysiological studies demonstrating that HlyA forms a water filled, cation-specific ion channel with a diameter of approximately 2 nm (26). Next to pore formation (4), Ca^2+^ oscillations at sub-lytic concentrations of the toxin (27), inflammasome activation (28), and cell death based on apoptotic, necrotic or pyroptotic processes (29, 30) have been described.

The nature of the HlyA receptor in the host’s plasma membrane has also not been fully resolved. In principle, two major pathways might exist, which both are supported by experimental data. First, a receptor-independent mode of action. This proposal is based on studies using artificial membranes (31, 32) and the observation that HlyA displays a non-saturable binding to rabbit erythrocytes (33). The identification of the heterodimeric integrin LFA-1 (α_L_ß_2_) as a receptor represented the starting point of a receptor-dependent mode of action. The receptor was later narrowed down to the ß_2_ subunit (CD18) (34, 35). A similar result was obtained for CyaA (36, 37), which recognized the integrin α_M_ß_2_ (CD11b/CD18). Interestingly, binding occurred to N-linked carbohydrates of the integrin (36). This would also explain why CyaA or HlyA can bind erythrocytes that lack LFA-1, but contain glycosylated proteins such as glycophorin, which was shown to function as a receptor for HlyA as well (38) and likely possesses a similar or identical glycosylation pattern.

In general, the primary structure of the secretion signal is not conserved among RTX proteins, the HlyA secretion system was shown to tolerate some non-native RTX toxins as substrates such as for example CyaA from *Bordetella pertussis*, FrpA from *Neisseria meningitidis*, PaxA from *Pasteurella aerogenes* or LktA from *Mannheimia haemolytica* (39–42). However, the secretion of these heterologous RTX proteins was not quantified or demonstrated to be efficient in terms of amounts of secreted protein. Furthermore, different experiments showed that the HlyA T1SS could secrete fusion proteins. The exchange of the HlyA secretion sequence with the C-terminus of LktA still facilitates secretion of HlyA (43). At the same time non-related, but slow-folding proteins carrying the HlyA secretion sequence are recognized and transported (13).

MbxA from *Moraxella bovis* shares a sequence identity of 42% with HlyA and is a key factor in the pathogenicity of *Moraxella bovis* (44, 45). This animal pathogen is the cause of the most prevalent ocular disease in cattle, the infectious bovine keratoconjunctivitis (IBK) (46, 47). Studies with supernatants of pathogenic *M. bovis* cultures revealed a cytotoxicity against bovine neutrophils, erythrocytes and corneal epithelial cells attributed to a possibly secreted cytotoxin (48–51). A classic RTX operon that comprises MbxA together with the putative acyltransferase MbxC along with the transporter components was later identified only in pathogenic, hemolytic *M. bovis* strains but not in non-hemolytic strains. With a length of 927 amino acids, MbxA is approximately 10% smaller than HlyA. Like for HlyA, sequence alignment and topology predictions revealed two potential acylation sites as well as an ambiguous membrane interaction domain at the N-terminal part of MbxA (44, 52, 53), while the GG- repeats are located closer to the C-terminus of MbxA compared to the setting in HlyA. Altogether, MbxA has a putative domain organization that is typical for pore-forming RTX toxins (Fig. 1).

A functional characterization of MbxA requires purified and active protein. So far, MbxA has only been produced for vaccination purposes as precipitated or partially solubilized protein (54) or as C-terminal fragments (55, 56).

Here, we established a complete surrogate T1SS in an *E. coli* laboratory strain by a combination of the UPEC HlyA T1SS with the heterologous substrate MbxA that secretes active MbxA or its precursor proMbxA. Based on this result, we developed a rapid and efficient purification procedure for both MbxA species from the supernatant of secreting *E. coli* cells. This approach is different from the cytosolic production and isolation of active RTX toxins from inclusion bodies such as for example CyaA from *Bordetella pertussis* (57) or LtxA from *Aggregatibacter actinomycetencomitans* (58). In our set-up, recombinant MbxA was efficiently activated by HlyC, but with an unexpected acylation pattern compared to HlyA. In contrast to the previously shown activity against bovine and ovine cells, we further observed that MbxA displays a host species independent cytotoxic activity against human epithelial cells and leukocytes. Using live-cell imaging, we finally demonstrate a rapid and distinct membrane blebbing phenotype in human epithelial cells in response to permeabilization by MbxA. RTX toxin induced plasma membrane blebbing has been so for only described for the multifunctional-autoprocessing RTX (MARTX) toxin RtxA1 from *Vibrio vulnificus* (*59*). This multidomain protein, composed of approximately 5200 amino acids, contains six putative effector domains including a pore-forming domain. Our results thus demonstrate that the ability of MbxA to induce membrane blebs is encoded in the pore-forming domain of this ‘classić RTX toxin and might be a general feature of this family of membrane damaging proteins.

## Results

### MbxA is secreted and activated by the HlyA system

To produce MbxA employing the UPEC-derived HlyA T1SS secretion system in *E. coli* BL21(DE3), the *mbxA* gene containing a hexa-histidine tag (*6H-*mbxA) for ease of purification was introduced into the pSU2726 vector (60) resulting in the plasmid pSU-*6H-mbxA*. The UPEC-derived HlyA T1SS inner membrane components HlyB and HlyD were expressed from the pK184-*hlyBD* vector, while endogenous BL21(DE3) TolC is chromosomally encoded (13). In a two-plasmid-system comprised of pK184-*hlyBD* together with pSU-*6H-mbxA* or pSU- *hlyC/6H-mbxA*, *mbxA* could either be expressed in the presence of the transporter components or additionally in the presence of the activating acyltransferase HlyC. *E. coli* BL21(DE3) that carry both, pK184-*hlyBD* and pSU-*hlyC/6H-mbxA,* showed hemolytic activity when grown on sheep blood agar, while in the absence of HlyC no halo formation occurred. Colonies harboring only pSU-*hlyC/6H-mbxA* did not cause hemolysis indicating that secretion via the T1SS and not cell lysis is not the reason for toxic activity of MbxA. This indicates that the secretion of active MbxA was indeed mediated by heterologous *E. coli* HlyBD-TolC (Fig. 2). The IPTG-induced expression of HlyBD together with MbxA only or MbxA with HlyC in liquid cultures led to an accumulation of an approximately 100 kDa protein, proMbxA or MbxA as verified by MS analysis (see below), respectively, in the medium (Fig. 3). In the absence of the HlyC protein, secretion levels of proMbxA were higher during 5 h of expression. While proMbxA was stable in the supernatant, MbxA tended to accumulate in the foam floating on the medium.

**Fig. 2.**
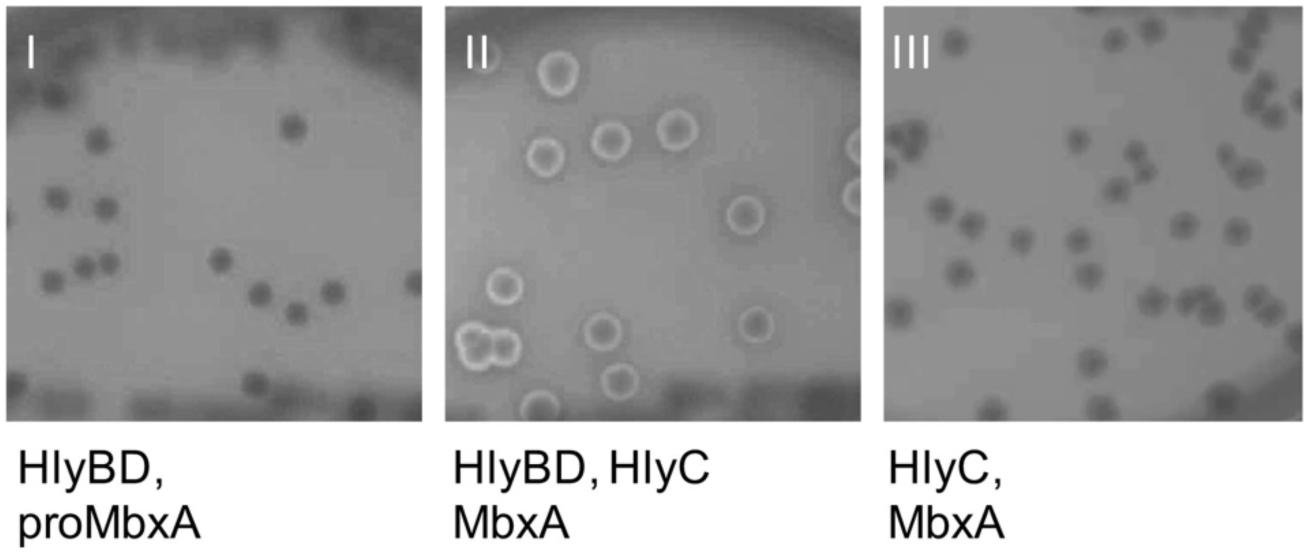
*E. coli* BL21(DE3) grown on Columbia agar with 5% sheep blood harboring the plasmids pK184-*hlyBD* with pSU-*6H-mbxA* (I), pK184-*hlyBD* with pSU-*hlyC*/*6H-mbxA* (II) or pSU-*hlyC*, *6H- mbxA* alone (III). Halo formation around colonies that expressed MbxA and HlyC together with the transporter components HlyBD (II) and absence of hemolysis around colonies that express only MbxA and HlyC (III) indicates functional secretion of active and acylated MbxA. Thus functional MbxA is not released from *E. coli* without additional expression of the HlyBD secretion system.

**Fig. 3.**
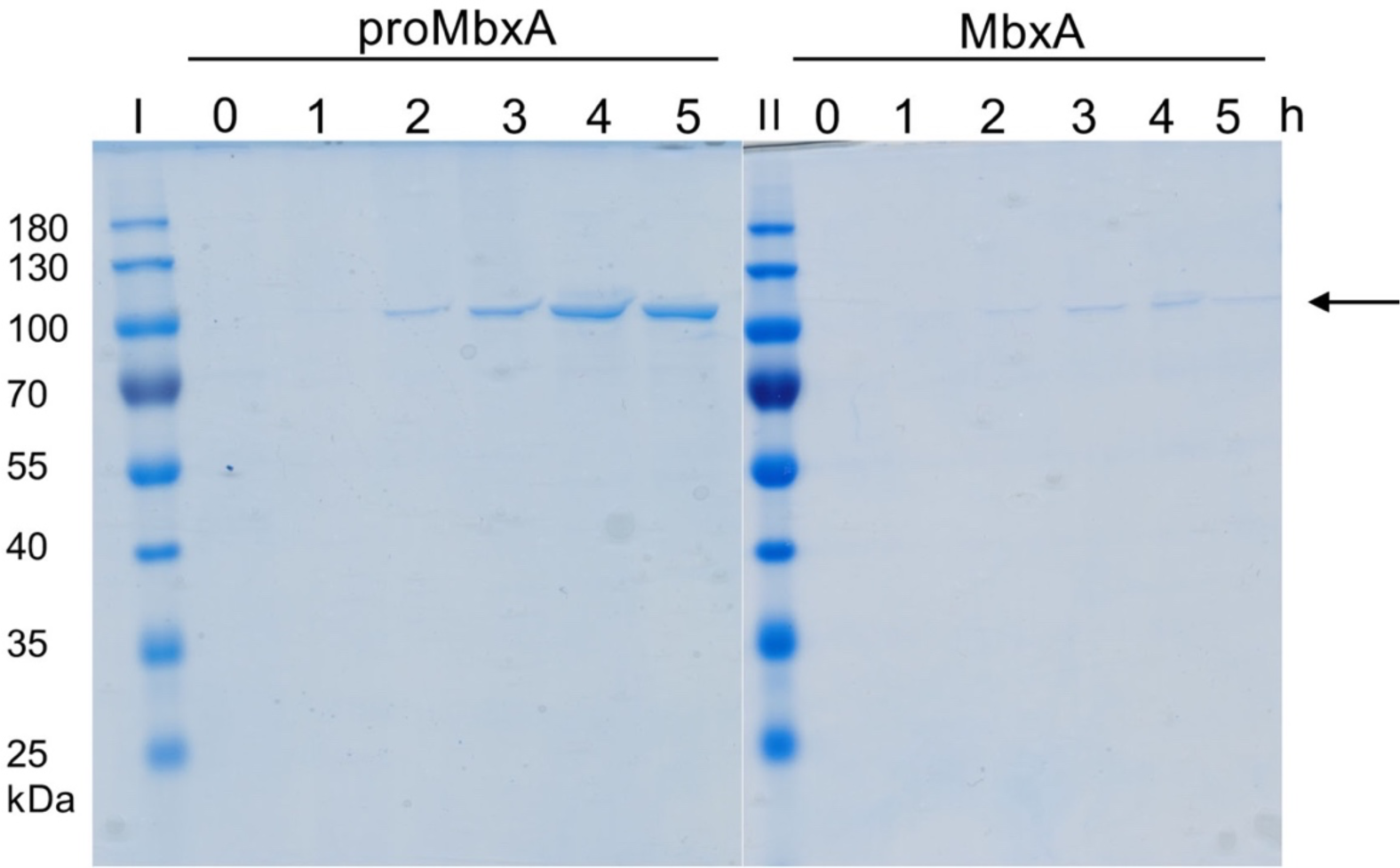
Heterologous secretion of proMbxA and MbxA via the HlyBD system. The Coomassie Brilliant Blue (CBB) SDS PAGE of culture supernatants of *E. coli* BL21(DE3) co-expressing HlyBD and MbxA (proMbxA, left panel) or HlyBD, HlyC and MbxA (MbxA, right panel) show accumulation of proMbxA or MbxA during a period of 0-5 h of expression (indicated by the numbers above the gel) after induction with IPTG. Samples of the supernatant were applied to the SDS gel without additional concentration by TCA precipitation. Molecular weight markers in kDa are shown in lanes I and II. An arrow indicates proMbxA and MbxA.

### Purification of proMbxA and MbxA from the supernatants of E. coli cultures

From the *E. coli* BL21(DE3) pK184-*hlyBD*, pSU-*6H-mbxA* system approximately 6 mg/L of precursor protein proMbxA was recovered from the supernatant using Ni^2+^ affinity chromatography. A subsequent size exclusion chromatography (SEC) and SDS PAGE analysis revealed that two separable species of proMbxA were present (Fig. 4). Determination of the molecular weight (MW) by multi angle light scattering (MALS) analysis confirmed that the first peak with an experimentally determined molecular weight (MW) of 201.5 ± 1.1 kDa corresponded to proMbxA dimers (theoretical MW 201.2 kDa). The second peak contained monomeric proMbxA as the determined MW of 93.9 ± 0.3 kDa fitted to the theoretical mass of 100.6 kDa (Fig. 5). The plasmid combination pK184-*hlyBD* and pSU-*hlyC/6H-mbxA* allowed the isolation of activated MbxA that was solubilized with 6 M urea from the foam of the expression culture and was afterwards refolded in the presence of Ca^2+^. It was equally purified via IMAC and subsequent SEC. MbxA eluted from the Superose 6 Increase column earlier (elution volume 13.6 ml) and in a broader peak than the proMbxA dimer (elution volume 14.9 ml), which is due to acylation and eventually indicates the presence of a higher oligomeric species of MbxA, probably including MbxA dimers and a small fraction of monomers (elution volume 17.1 ml) (Fig. 4). CBB staining confirmed the purity of proMbxA and activated MbxA already after IMAC (Fig. 4b).

**Fig. 4.**
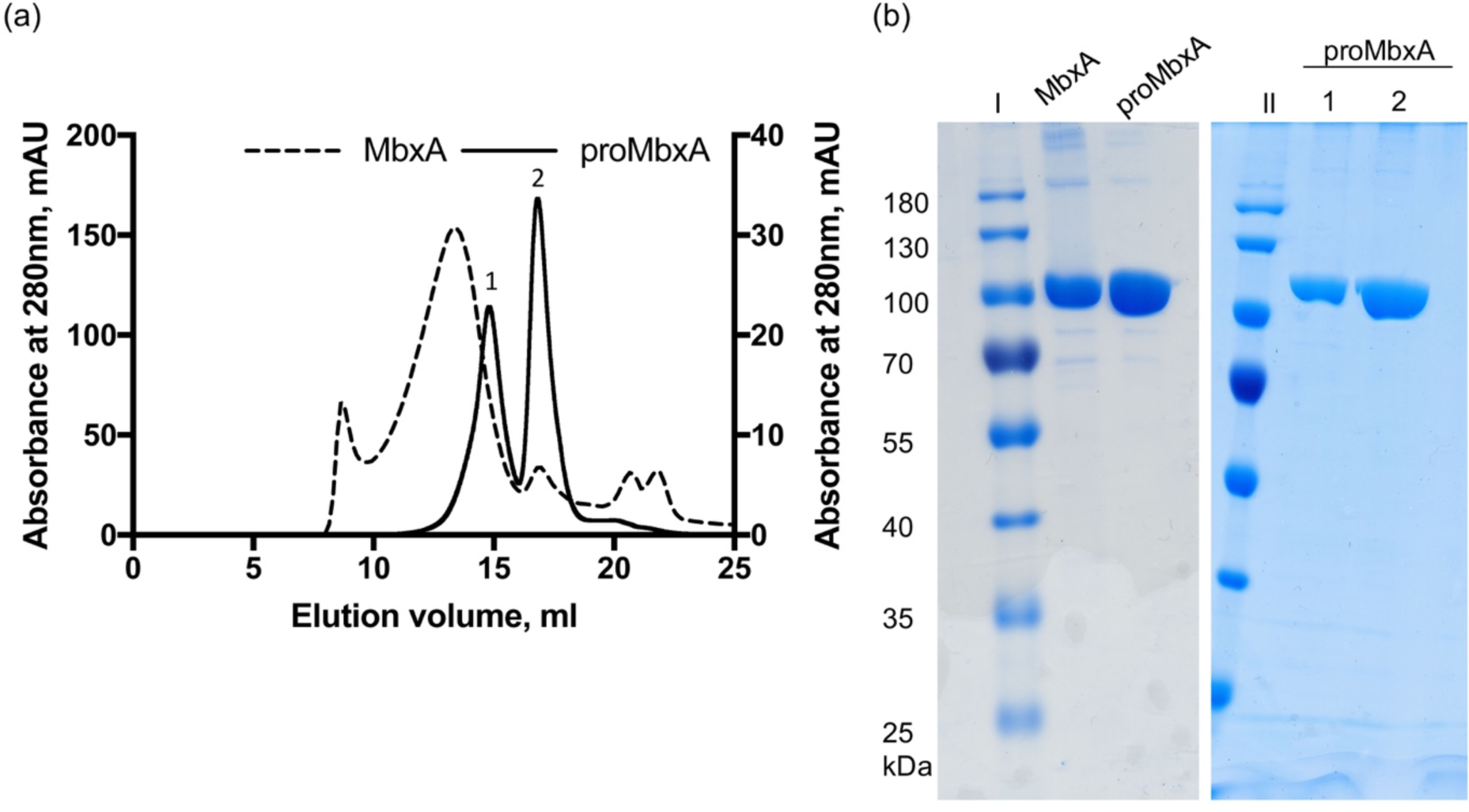
(a) SEC chromatographs of samples of active MbxA (dashed line) and proMbxA (solid line), respectively, applied to a Superose 6 Increase 6/30 column (GE Healthcare). The absorbance at 280 nm of the elution profile of proMbxA was plotted on the left Y axis, the elution profile of active MbxA on the right Y axis. proMbxA eluted in two separate main peaks, marked 1 and 2 in the chromatogram. (b) CBB stained SDS PAGE gels of pooled fractions of active MbxA and proMbxA obtained from IMAC purification (left panel) and used for SEC. proMbxA was separated in two species, peak 1 and peak 2, during SEC and evaluated by SDS PAGE (right panel). Molecular weight markers in kDa are shown in lanes I and II.

**Fig. 5.**
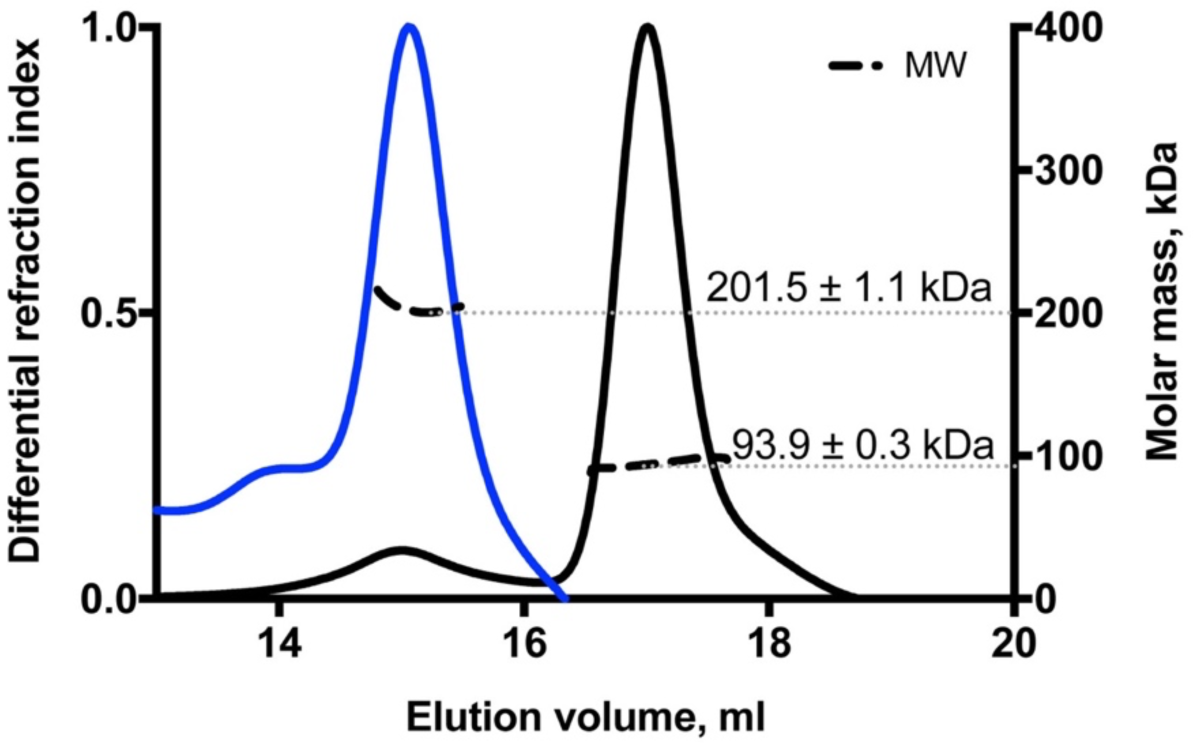
MALS coupled to SEC analysis of the two proMbxA species separated via SEC (blue line peak 1 of Figure 4, black line peak 2 of Figure 4). The calculated molecular weight of 201.5 ± 1.1 kDa of the first peak (blue) corresponds to a dimer of proMbxA (theoretical MW (6H- MbxA)_2_= 201.2 kDa). The molecular weight of 93.9 ± 0.3 kDa of the second peak (black) confirms a monomeric species of proMbxA (theoretical MW 6H-MbxA= 100.6 kDa).

### Active MbxA is acylated by HlyC at two conserved acylation sites

Besides HlyA, several other pore-forming RTX toxins are known to require a post-translational acylation at two or, as in the example of CyaA, at least one conserved internal lysine residue to confer toxic activity (20, 61–65). For HlyA, acylation of both acylation sites was shown to be required for toxicity (20). The amide linked fatty-acylation is facilitated by an acyltransferase encoded upstream of the RTX gene and an acyl carrier protein (66). The presence of the *mbxC* gene in the *M. bovis* operon indicates that expressed MbxA is likely acylated (44). The lysine residues K536 and K660 of MbxA correspond to the acylation sites at K564 and K690 of HlyA (Fig. S1) (20, 44). The areas around the two acylation sites, residues H516-G566 and K640- Q680, share a sequence identity of approximately 46% with those of HlyA. This identity is comparable to the overall identity of both proteins. To analyze a possible acylation status of proMbxA expressed in the absence of HlyC and MbxA co-expressed with HlyC, we used mass spectrometry. No acylation was detected for proMbxA in the absence of HlyC. In contrast, in the presence of HlyC MbxA peptides containing acylated lysine residues were identified at the predicted sites, K536 and K660, respectively. No peptide with non-acylated K536 or K660 was detected in the acylated MbxA variant. Myristoylation (C_14_ acylation) as well as hydroxy myristoylation (C_14_-OH*) were determined for all three detected peptides that covered the first acylation site K536 (ERLTNG**K**YSYINK, LTNG**K**YSYINK and LTNG**K**YSYINKLK) (Fig. 6 and Tab. S1). Each of those peptides was identified by more than one fragment spectrum recorded by the mass spectrometer resulting in a total of 24 informative spectra for C_14_ modification with additional 5 informative spectra for C_14_-OH* modifications of these three peptides. Those peptide spectrum matches (PSM) can be used as semi-quantitative measure for the occurrence of peptides and peptide modifications. The peptide LTNG**K**YSYINK was additionally found to be modified with C_13_, C_15_ and C_16_ acylations as well as a C_12_ hydroxy acylation. These modifications with C_13_, C_15_, C_16_ and C_12_-OH* fatty acids are probably less frequent as they were detected only with 2, 4, 1 and 1 PSM, respectively. The second acylation site, K660, was only found to be acylated at a low level with C_14_ fatty acids (Fig. 6a). As no other fatty-acylated lysine residues were observed, we conclude that modification of MbxA with the non-native, but promiscuous acyltransferase HlyC, here referred to as cross-acylation, targets only the predicted acylation sites albeit to different degrees (Tab S1).

**Fig. 6.**
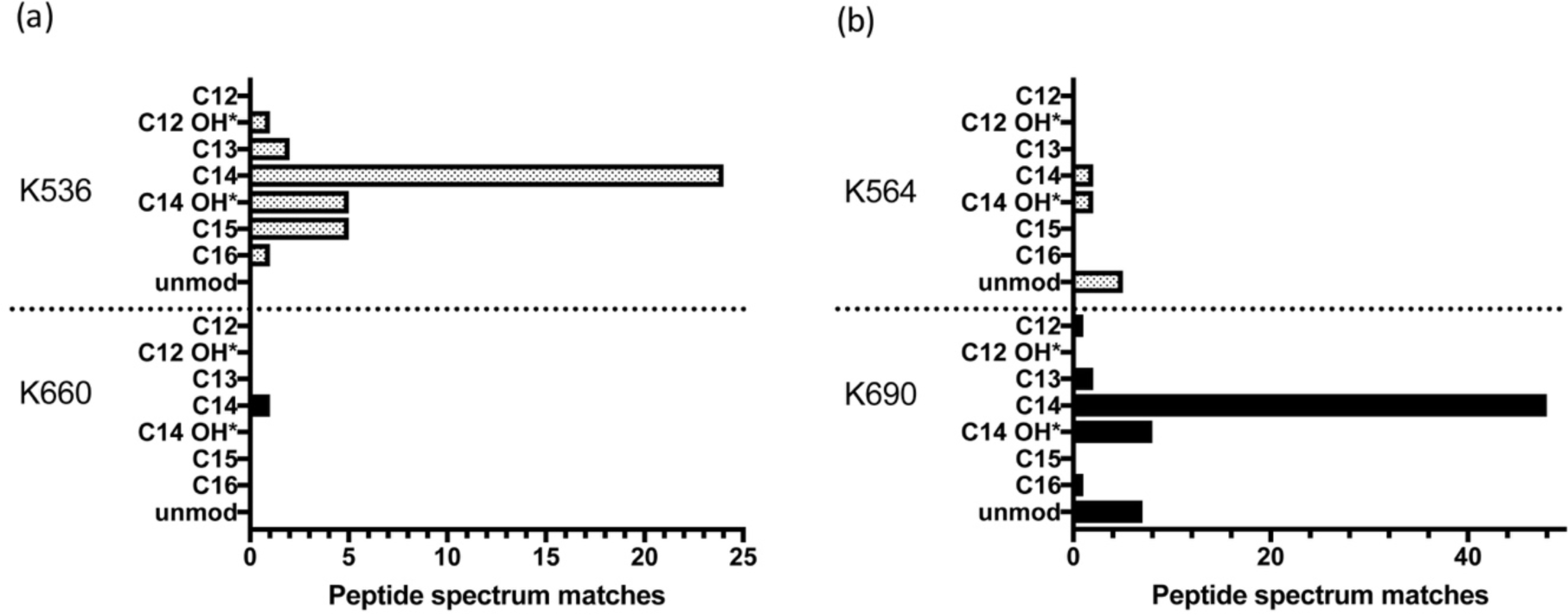
MS analysis of MbxA acylation (a) and HlyA acylation (b). MbxA and HlyA were *in vivo* acylated by co-expressing HlyC (67). For both acylation sites in MbxA and HlyA, K536 and K660 or K564 and K690 respectively, all detected acyl modifications ranging from C_12_ to C_16_ fatty acids including hydroxy fatty acids are shown with the corresponding number of recorded PSM. Mass shifts contributing to hydroxy fatty acids could possibly originate from the oxidation of tyrosine residues in the peptide fragment and PSM of supposed hydroxy acylations are therefore marked (*). For proMbxA and proHlyA, both expressed in the absence of HlyC, only unmodified peptides were detected. Acyl modifications were detected only in peptides covering the predicted lysine residues K536 and K660 of MbxA. The PSM of C_14_ modified sequences suggest that the predominant modification is myristoylation.

To test whether HlyC confers the same modifications to MbxA as to HlyA in the *E. coli* BL21(DE3) background, proHlyA (absence of HlyC) and HlyA (presence of HlyC) were analyzed for their acylation status. For HlyA, peptides acetylated at both known acylation sites (20) could be identified (Tab S1). At K564 peptides modified with C_14_ acylation and C_14_-OH* acylation contributed both with 2 PSM. The second acylation site, K690, was predominantly found acylated with C_14_ fatty acids (48 PSM) but also hydroxy acylated with C_14_-OH* modifications (8 PSM). Moreover, and in contrast to the second acylation site of MbxA, also less frequent modifications, namely C_12_, C_13_ and C_16_ acylations were detected at the K690 acylation site of HlyA with 1, 2 and 1 PSM, respectively. Additionally, unmodified peptides containing K564 and K690 were detected for HlyA (5 and 7 PSM, respectively) (Fig. 6b). Based on the PSM data, we assume that the vast majority of HlyA proteins are acylated, as in proHlyA no acylated peptides were determined, but unmodified peptide variants including K564 and K690 with 114 and 68 PSM, respectively. Interestingly, the ratio of acylation of the two sites is invers. While in MbxA the first site is more efficiently acylated, the second site showed a more efficient acylation in HlyA, although the same acyltransferase catalyzed the reaction in the context of the laboratory strain BL21(DE3). The presence of peptides resulting from cleavage directly after the less frequently detected second acylation site of MbxA and the first acylation site of HlyA further indicates that these sites were not fully acylated as a modification would have masked the cleavage site (Tab. S1).

The analysis of the *in vivo* acylation of HlyA by HlyC confirms the acylation of the K564 and K690 acylation sites. Similar to K536 in MbxA, C_14_ and C_14_-OH* acylations account for the majority of PSM of acyl modified peptides at K690 in HlyA.

### MbxA is cytotoxic to human epithelial cells and T cells

As acylated MbxA displayed hemolytic activity on sheep blood agar, human epithelial cells (HEp-2) and human T lymphocytes (Jurkat) were tested for their susceptibility to MbxA with a lactate dehydrogenase (LDH) release assay. Both cell lines were susceptible to MbxA-induced LDH release, which implies cell membrane damage, and showed a sigmoidal dose-response curve with increasing MbxA concentrations (Fig. 7). This is indeed a novel finding since so far only bovine cells were reported to be targeted by MbxA. The half-maximal effect (cytotoxic dose 50, CD_50_) of MbxA against HEp-2 cells occurred at a concentration of 28.1 ± 4.7 nM. For Jurkat cells 50% cytotoxicity was reached at a lower concentration of MbxA resulting in a CD_50_ of 17.7 ± 3.9 nM. Importantly, proMbxA displayed no cytotoxic effect after 1 h. The calculation of the CD_50_ values was performed under the assumption that the entire pool of MbxA is active. Thus, the values reported likely correspond to a upper limit for the CD_50_ values. These data demonstrate that MbxA is toxic to both, human epithelial and immune cells, in a similar concentration range, but strictly requires activation by acylation to confer membrane-damaging activity.

**Fig. 7.**
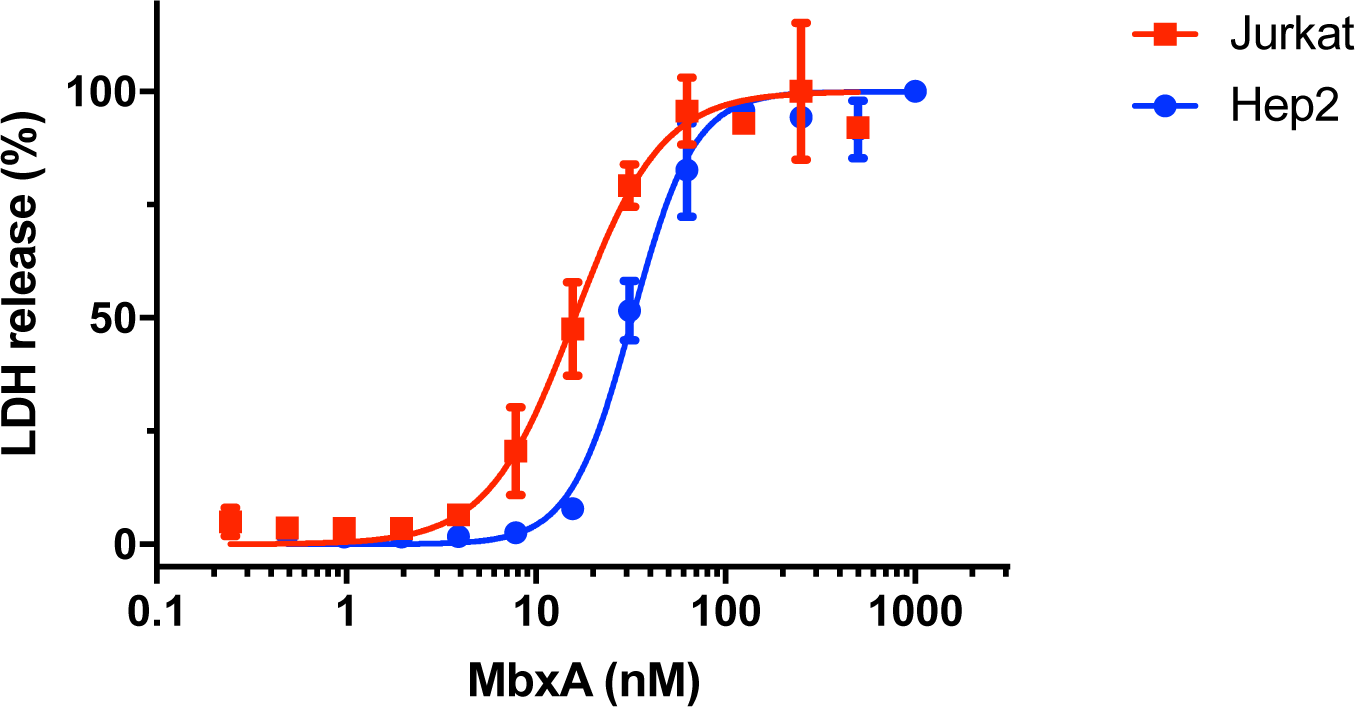
MbxA induces LDH release in human epithelial cells (HEp-2, 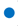) and human T cells (Jurkat, 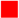), respectively. The cytotoxicity mediated by MbxA-induced membrane damage was measured using an LDH release assay. LDH release into the supernatant was measured after 1 h of incubation with MbxA. For the CD_50_ determination the MbxA cytotoxicity was plotted against the MbxA concentration and fitted with GraphPad Prism 7 according to equation 2 (see materials and methods). For each cell type, the highest LDH release was set to 100%. For simplicity, measurements with proMbxA were not included as no LDH release, even at concentrations of 1 µM, was detected.

### proMbxA cannot protect human cells from MbxA-induced membrane damage

As proMbxA did not display any cytotoxicity against epithelial cells or T cells we further proceeded to assess if proMbxA can protect any potential MbxA binding sites by binding and shielding the recognition site on the surface of human cells. Preincubation with 250 nM of proMbxA for 30 min at 37 °C before addition of a serial dilution of active MbxA to HEp-2 cells and Jurkat cells had no effect on the dose-response curve and the CD_50_ value compared to MbxA alone (Fig. 8). The presence of proMbxA did not decrease the sensitivity of any of the two cell lines indicating that proMbxA cannot interfere with the cytotoxic action of MbxA.

**Fig. 8.**
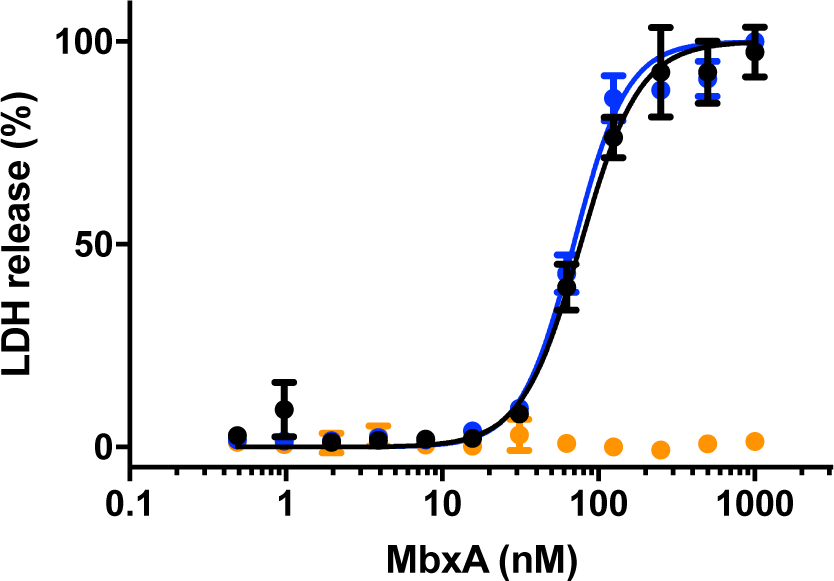
proMbxA does not protect epithelial cells (HEp-2) from MbxA-induced membrane damage. Preincubation with 250 nM of proMbxA for 30 min at 37 °C resulted in a nearly identical dose-response curve to MbxA (black) compared to treatment with MbxA alone (37 °C for 30 min, blue). proMbxA was not cytotoxic (orange). The cytotoxicity mediated by MbxA- induced membrane damage was measured using an LDH release assay. LDH release into the supernatant was measured after 1 h of incubation with MbxA. The relative LDH release was calculated from the maximal LDH release reached in each measurement. For the CD_50_ determination the MbxA cytotoxicity was plotted against the MbxA concentration and fitted with GraphPad Prism 7 according to equation 2 (see material and methods).

### MbxA induces membrane permeabilization and membrane blebbing in HEp-2 cells

To study the dynamics and the appearance of the changes in the membrane morphology of HEp-2 cells, which occurred upon incubation with MbxA in more detail, we used confocal live-cell imaging. Therefore, the plasma membrane of HEp-2 cells was stained by CellMask™ Deep Red prior to incubation with MbxA. In parallel, the membrane impermeable high affinity DNA dye Sytox® Green was supplied in the surrounding medium of the HEp-2 cells, to monitor the permeabilization of the HEp-2 cells indicated by an increasing Sytox® Green signal. Different final concentrations of MbxA (250 nM, 30 nM and 10 nM) and 250 nM of proMbxA were provided to the HEp-2 cells and confocal micrographs were acquired every 15 seconds for a total of 20 minutes. When HEp-2 cells were treated with 250 nM MbxA (Fig.9), Sytox® Green enters the nuclei rapidly within the first minutes and stains the DNA with increasing intensity during the observation period of 20 minutes. Immediate alteration of the plasma membrane morphology could be observed as deformation and blebbing on the cell boundaries (second row) and prominent occurrence of large membrane blebs was seen above the HEp-2 cell layer (third row). To further verify and observe the dose-dependent effect of MbxA in live-cell experiments, Sytox® Green influx was analyzed when HEp-2 cells were challenged with 30 nM of MbxA (a concentration around the CD_50_ determined by the LDH release assay; Table 2) and 10 nM (a concentration that resulted in minimal effects; Fig. 7).

**Fig. 9.**
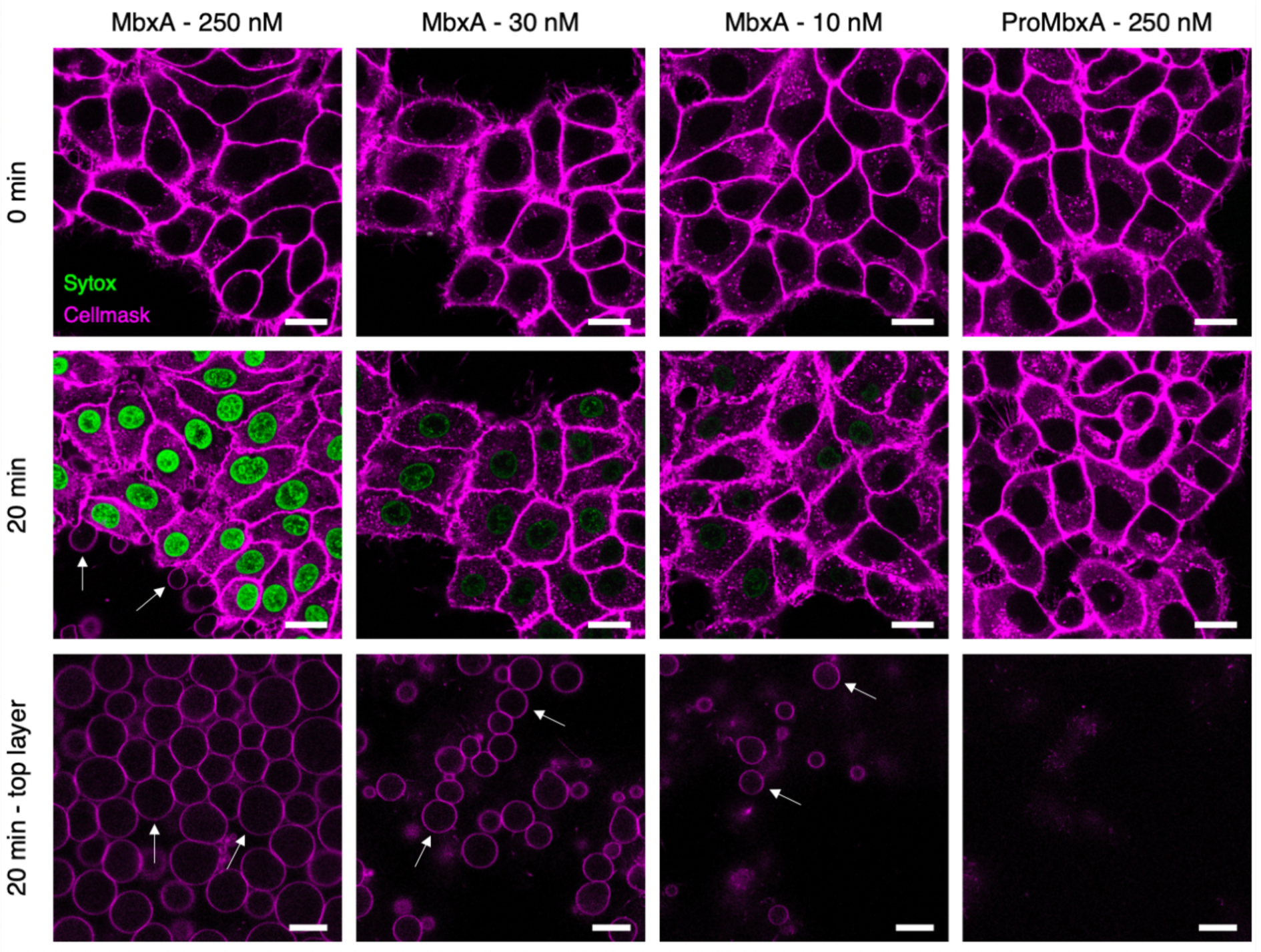
Confocal microscopy images of MbxA-induced membrane permeabilization and membrane blebbing in HEp-2 cells. HEp-2 cells were incubated with MbxA (250 nM, 30 nM and 10 nM) or proMbxA (250 nM) and analyzed at time point zero (first row) and after 20 min (second row). Plasma membrane was stained by CellMask™ Deep Red and membrane permeabilization was monitored using Sytox® Green staining of the nuclear DNA. Incubation of HEp-2 cells with 250 nM, 30 nM and 10 nM MbxA resulted in DNA staining with Sytox® Green cells in decreasing intensity corresponding to the decreasing concentration of MbxA and hence in membrane permeabilization. DNA staining was not observed when HEp-2 cells were incubated with 250 nM proMbxA. Membrane blebs were observed in the same focus layer as the HEp-2 cells (second row; white arrows) and above the HEp-2 cells (third row; white arrows). While incubation with 250 nM and 30 nM MbxA led to severe appearance of membrane blebbing, incubation with 10 nM of MbxA was less noticeable and 250 nM proMbxA incubation did not lead to any formation of blebs. Scale bar: 20 µm.

To understand the correlation between membrane permeabilization and cell death using different MbxA concentrations, quantification of Sytox® Green signal was performed per nucleus over time (Fig. 10). Incubating HEp-2 cells with 30 nM MbxA also resulted in a detectable Sytox® Green staining of the DNA, but the slope of the intensity per nucleus over time was notably lower compared to the Sytox® Green staining observed with 250 nM MbxA (Fig. 9 and 10). Additionally, membrane blebbing of HEp-2 cells with 30 nM MbxA occurred slower and formation of blebs was less severe compared to 250 nM MbxA. Using only 10 nM MbxA during the measurement, even less or occasionally no Sytox® Green intensity per nucleus was detected, and a few membrane blebbing events were observed at the cells and above the cell layer (Fig. 9). While MbxA incubation of HEp-2 cells leads to membrane permeabilization and membrane blebbing, incubation of HEp-2 cells with 250 nM proMbxA resulted in neither Sytox® Green staining of the DNA nor observable changes in membrane morphology such as blebbing, which is indicative of a preserved plasma membrane integrity (Fig. 9 & 10). Similar results were observed using AlexaFluor488-labeled Wheat Germ Agglutinin (WGA-488) as membrane marker and propidium iodide (PI) staining for nuclear DNA as measure for permeabilization (Fig. S4 & S6).

**Fig. 10.**
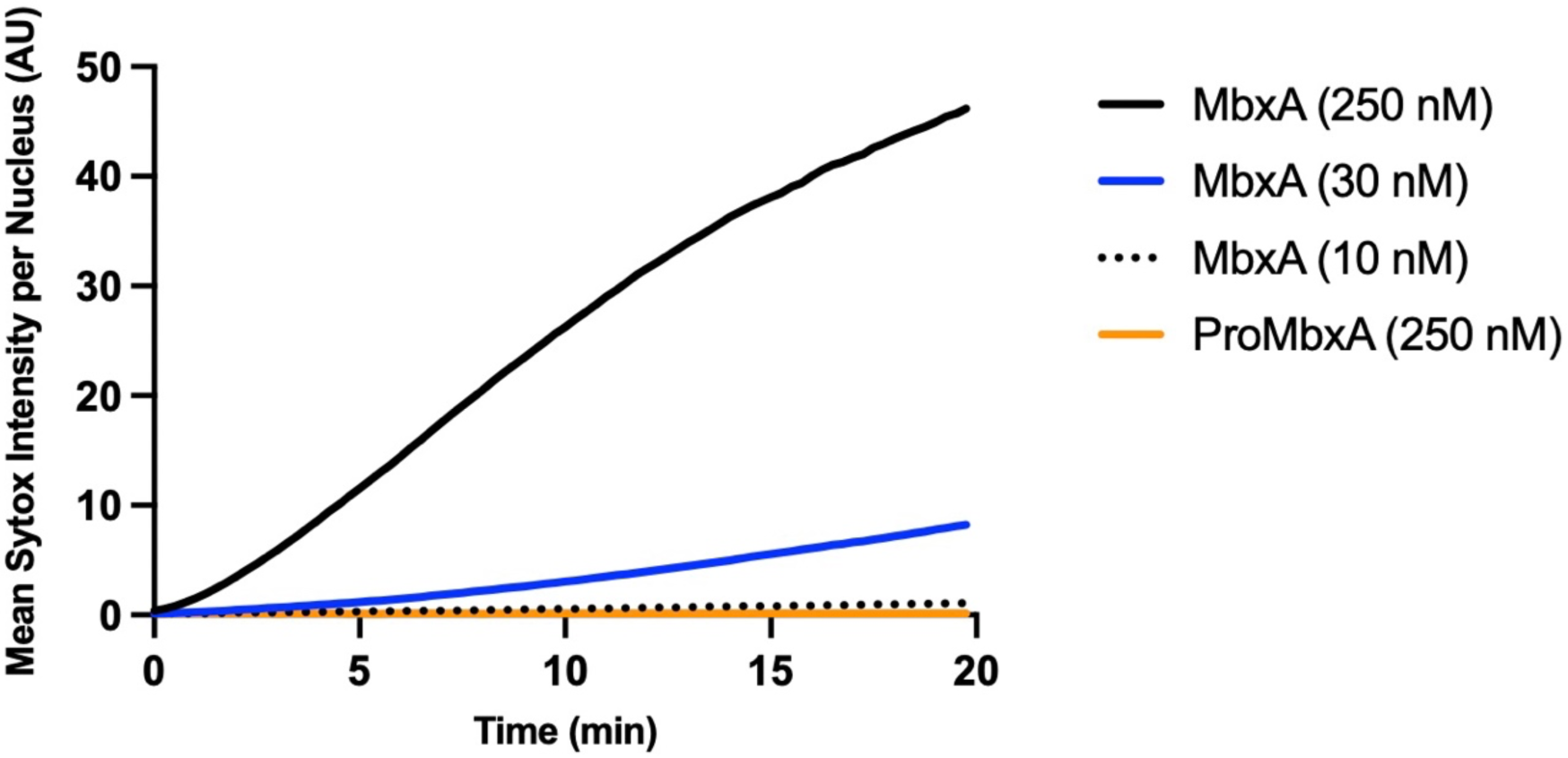
Quantification of MbxA-induced permeabilization of HEp-2 cells. Mean Sytox® Green intensity per nucleus of HEp-2 cells (y-axis) was continuously measured over a period of 20 minutes (see mateirals and methods for details) and is plotted against time (x-axis) for different MbxA concentrations (250 nM, 30 nM and 10 nM) and 250 nM. Sytox® Green staining of the HEp-2 cellś DNA is faster and more intense when 250 nM MbxA were used and is reduced at lower MbxA concentrations (30 nM and 10 nM). In contrast to that, no significant Sytox® Green staining of the nuclear DNA was observed with 250 nM proMbxA. (Number of nuclei used for quantification: 250 nM MbxA – 71, 30 nM MbxA – 74, 10 nM MbxA – 79, 250 nM proMbxA – 62). Nuclei have been analyzed from two independent experiments with two biological replicates each.

Together, this data imply that HEp-2 cells were almost immediately permeable for Sytox® Green after contact with MbxA in a dose-dependent manner and underwent changes in membrane morphology like blebbing caused or accompanied by cell death. Moreover, these observed effects must be a functional trait of active MbxA, since the inactive precursor proMbxA did not induce permeabilization, morphological changes or cell death.

### Labeled MbxA localizes to the plasma membrane of HEp-2 cells

In order to visualize the localization of MbxA, MbxA was purified and labeled using Atto-488 at the N-terminus. HEp-2 cells were incubated with 250 nM Atto488-MbxA^S9C^ for 30 min.

Before a washing step with fresh medium, Atto488-MbxA^S9C^ was clearly detectable in the medium (Fig. 11a), while induced morphological changes of the plasma membrane were observed as membrane blebs (Fig. 11b). The strong fluorescence dots in Fig. 11a likely represent aggregated protein as they were hardly detectable after the washing step. To analyze the localization of Atto488-MbxA^S9C^ after unbound protein in the cell supernatant was removed and replaced by fresh media, cells were imaged again (Fig. 11d). Accumulations of Atto488-MbxA^S9C^ tracing the shape of the HEp-2 cells were clearly visible when the unbound Atto488-MbxA^S9C^ was removed as shown in the overview images (Fig. 11d) and in the super– resolution micrographs acquired using the Airyscan technique (Fig. 11e). This visualizes the localization of MbxA to the plasma membrane of HEp2 cells, which goes together with permeabilization, potential pore formation and subsequent cell death. Incubation with free Atto488 as a control did not lead to accumulation of the dye in the membrane (Fig. 11c before washing and Fig. 11f after washing) underlining the specific insertion of MbxA into the membrane.

**Fig. 11.**
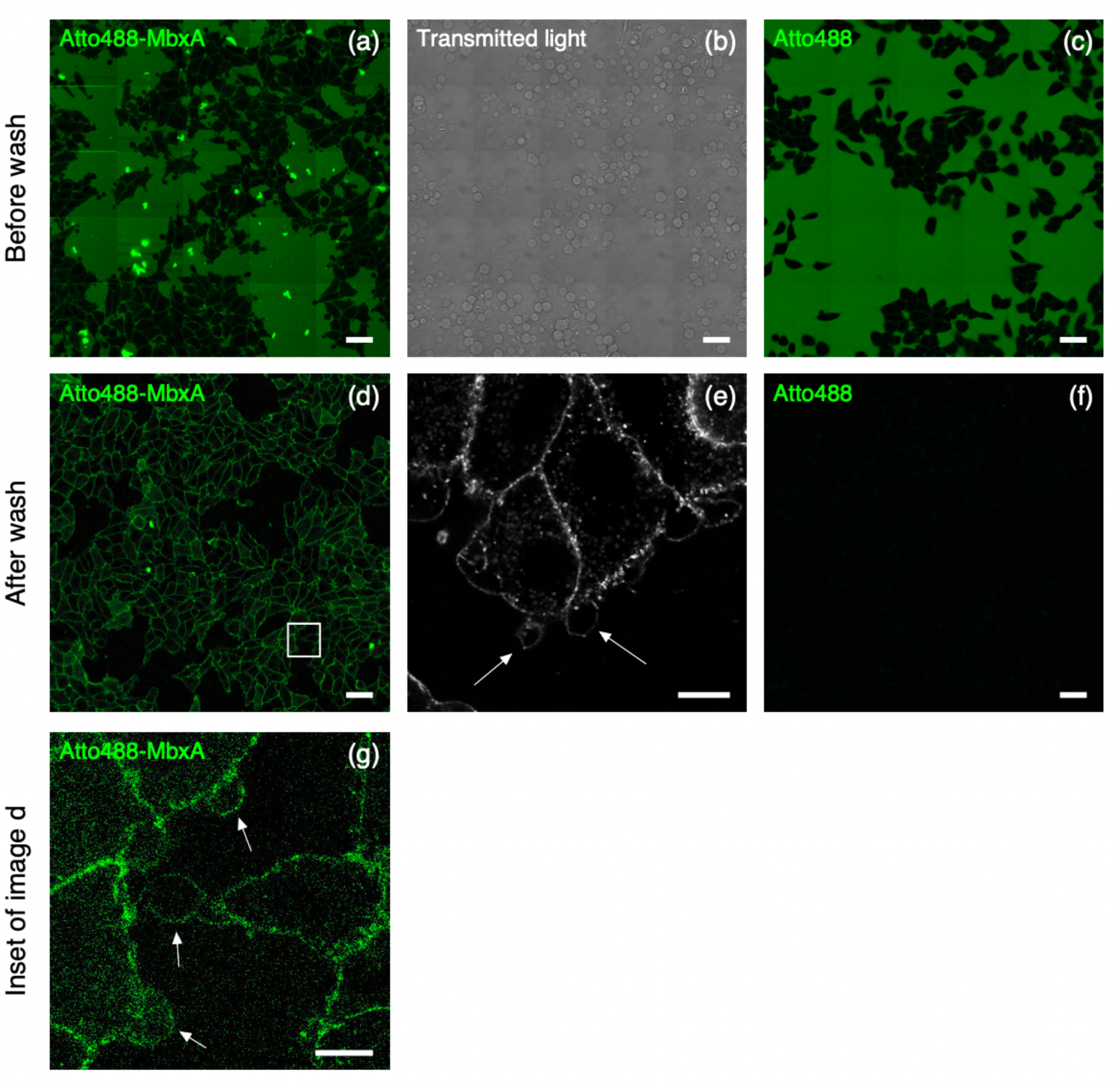
Incubation experiment of Atto488-MbxA^S9C^ with HEp-2 cells. HEp-2 cells after 30 min of incubation with 250 nM Atto488-MbxA^S9C^, before (a and b) and after (d and e) removing excess unbound Atto488-MbxA^S9C^. Confocal overview images of fluorescent signal are shown in (a) and (d), transmitted light showing formation of blebbing process in (a) and (b) super-resolution Airyscan micrograph in (e). Accumulations of Atto488-MbxA^S9C^ tracing the outline of HEp-2 cells are clearly visible after removing the unbound Atto488-MbxA^S9C^ (c and d). Control experiment with Atto488 (dye used to label MbxA) is shown in (c) and (f). HEp-2 cells after 30 minutes of incubation with Atto488, before (c) and after (f) washing off the unbound Atto488 from surrounding medium. The control experiment confirmed that accumulations tracing the outline of HEp-2 cells in (d) and (e) are Atto488-MbxA^S9C^, not the free dye. White arrows indicate the position of membrane blebs. Scale bars: a, b, c, d, f: 50 µm; e: 10 µm.

## Discussion

MbxA is a cytotoxin of the bovine pathogen *M. bovis* and belongs to the RTX protein family (44). As an RTX toxin, MbxA is characterized by five conserved Ca^2+^ binding repeats that form the RTX domain and is likely secreted via a cognate T1SS. The necessary tripartite secretion system secreting MbxA is encoded together with an activating acyltransferase and the toxin gene on a pathogenicity island in the *M. bovis* genome (68). The presence of an intact RTX operon and cytotoxicity of *M. bovis* culture supernatant allows the conclusion that the MbxA secretion system and the MbxA toxin itself are functional (48, 49, 69). However, a detailed molecular analysis is still lacking.

Here we employed the secretion system of HlyA from the uropathogenic *E. coli* strain UTI89, considered as a prototype of RTX proteins, to secrete MbxA, thus creating a heterologous T1SS in *E. coli* BL21(DE3). In our recombinantly assembled secretion system the inner membrane components HlyB and HlyD (from UPEC *E. coli*) successfully recognize and interact with the secretion signal of MbxA (from *M. bovis*). The C-terminal secretion signals of RTX proteins do not harbor a conserved sequence identifiable by sequence analysis, but are likely recognized by a secondary structure element, presumably an amphipathic helix motif (70). This suggests that MbxA carries this required information likewise in its C-terminus as it is efficiently transported by the HlyA system and can be purified to homogeneity in amount sufficient for any biochemical, biophysical or cellular study. As a putative pore-forming toxin of the RTX family MbxA requires a post-translational activation by acylation of two internal lysine residues. Therefore, we completed the recombinant secretion system by expression of *E. coli* UTI89 HlyC instead of the putative *M. bovis* acyltransferase MbxC resulting in an hemolytically active *E. coli* BL21(DE3) strain (Fig. 2). Both, acylated MbxA and the precursor proMbxA, although in higher amounts, were secreted and facilitated purification from culture supernatants (Fig. 3). The difference in secretion efficiency is similar to proHlyA which is more efficiently secreted than its acylated version, HlyA (67). Our SEC analysis revealed that proMbxA exists as a dimer and a monomer, while the acylated MbxA forms higher oligomeric species, which presumably include dimers (Fig. 4). Similarly, proHlyA forms dimers when purified, but active HlyA is prone to aggregation (67). On the contrary, acylation of proCyaA results in formation of the active, monomeric CyaA species (57).

Mass spectrometric analysis of HlyC-acylated MbxA demonstrated that HlyC acylates the proposed lysine residues, K536 and K660, that correspond to the two acylation sites of HlyA (K564, K690) (20, 44). HlyC displays a clear preference for myristic acid (C_14_ fatty acids) and only small amounts of C_13_, C_15_ or C_16_ fatty acids appear at the first acylation site. Additionally potential C_12_-OH* and C_14_-OH* hydroxy acylations were detected at the first acylation site. The second acylation site is only found to be myristoylated (Fig. 6). We observed a similar preference for myristoylation and a similar variety of fatty acids utilized for modification of HlyA including C_12_, C_13_, C_14_, C_14_-OH* and C_16_ acyl chains. In contrast to MbxA, where preferentially the first acylation site is modified with different fatty acids, in the case of HlyA variable modifications are detected mainly at the second acylation site. Previous studies already showed that HlyA is predominantly myristoylated but reported a substantial percentage of C_15_ and C_17_ modifications of up to 32% (22), which we did not observe in HlyA expressed and acylated *E. coli* BL21(DE3). Another recent study underlined the preference of HlyC for C_14_ and C_14_-OH fatty acids and reported that the acylation pattern and choice of acylation sites is inherent to the acyltransferase in the case of non-secreted substrates (23). At the same time, our study shows that the acylation is not solely controlled by the acyltransferase as we observed that HlyC acylates the first acylation site with a variety of fatty acids in MbxA, while for HlyA more variation of the modification is found at the second site. This illustrates that the same acyltransferase, HlyC, displays different acylation patterns depending on the substrate protein. Interestingly, Osickova *et al*. (23) reported that mono- acylation of the second acylation site was also observed for the active species, while our study and the study of Stanley *et al*. (20) demonstrated that the double acylated version represents the active species. Assuming that the relative efficiency of acylation partially depends on the amount of the RTX protein present in the bacterial cytosol prior to secretion, the unmodified peptide fragments of HlyA detected in our system reflect its higher expression levels compared to MbxA. A very high level expression of the RTX protein that results in inclusion body formation as given in the study of Osickova et al. (23) may explain the incomplete modification of the first acylation site. Altogether, the HlyA system can successfully replace the complete T1SS of another RTX substrate and facilitate its recombinant activation and secretion in *E. coli*. This acylation is obviously the prerequisite for the toxic activity of MbxA. In contrast to previous studies focusing on the activity of MbxA against bovine cells (49–51, 71) we demonstrated cytotoxicity of MbxA outside of its known target species. Besides the hemolytic activity towards sheep erythrocytes, MbxA exhibited cytotoxic activity against human epithelial cells (HEp-2) and human T cells (Jurkat) as demonstrated by LDH release assays (Fig. 7) suggesting that MbxA might act species-independent. Importantly, proMbxA did no induce any LDH release indicating that acylation of MbxA was strictly required for its cytotoxicity. Additionally, preincubation with unmodified proMbxA was not able to block interaction of active MbxA with epithelial cells and T cells or hamper MbxA-induced membrane damage (Fig. 8). This suggests that without the fatty acid-acylation proMbxA cannot successfully bind potential receptors or surface structures on the human cells. This confirms that MbxA as a representative of RTX pore-forming toxins relies on posttranslational acylation. The necessity of acylation for cytotoxicity was previously shown for different RTX toxins and is linked to oligomerization, irreversible insertion into the membrane and / or an effective binding to a host cell receptor (64, 65, 72–74). At the same time, acylation is not required for binding to and insertion into membranes or pore-formation, at least for HlyA (60, 75, 76).

The cytotoxicity of MbxA, a toxin from a bovine pathogen, against human epithelial cell or leukocytes was, with a CD_50_ value of 28.1 ± 4.7 nM or 17.7 ± 3.9 nM (Fig. 7), similar to the activity reported for RTX toxins of human pathogens underlining that its activity is not confined to specific structures of bovine cells. HlyA for example is active at CD_50_ values ranging from approximately 0.2 nM to 30 nM depending on the cell line (77). The pore-forming activity of MbxA was previously described for bovine erythrocytes, in which MbxA induces leakage of K_+_ ions, cell swelling and finally lysis by forming pores of an estimated size of 0.9 nm (48). We performed live-cell imaging and super-resolution microscopy of HEp-2 cells challenged with MbxA and observed specific localization of MbxA to the plasma membrane and a distinct change in the membrane morphology. Besides permeabilization of the membrane indicated by PI and Sytox Green uptake, large spherical membrane protrusions of up to 20 µm diameter were induced by MbxA at concentrations ranging from 10 nM to 250 nM, but importantly not by proMbxA (Fig. 9). The formation of these protrusions is reminiscent of membrane blebbing, a phenomenon that is a part of apoptosis and necrosis (78, 79), but is also observed in healthy cells during cell division, spreading and migration (80–82). To the best of our knowledge, RTX toxin-induced formation of membrane blebs was only described for contact-dependent RtxA1 from *V. vulnificus* (*59*). In other words, the cytotoxic action of RtxA1 requires a physical contact of the bacterial cell with the host cell. Only then, a time-dependent expression of the protein was detected. In addition, RtxA1 is a multidomain protein and belongs the sub-family of MARTX toxins (83). Next to membrane blebbing, RtxA1 induced also cell rounding as a consequence of a re-arrangement of the actin cytoskeleton (59). In contrast to MbxA, RtxA1 contains not only a pore-forming domain but also four additional functional domains, one or more of which are responsible for the changes in cell rounding. For MbxA, which lacks this multidomain architecture, such cell rounding was hardly observed in our image analysis . This clearly suggests that the formation of membrane blebs is an intrinsic function of the pore-forming unit of MbxA and does not require any cofactor or cell-cell contact as in the case of RtxA1.

Membrane blebs form either when an increased intracellular pressure leads to tearing of the membrane from the cytoskeleton or when the actin cortex itself breaks and the excess membrane subsequently inflates (84–86). The lesions induced by MbxA probably might lead to an increase in the cell pressure by influx of medium components such as Ca^2+^ along the concentration gradient, which than leads to inflation of the excess membrane forming the bleb-like structure. In contrast to blebbing in healthy cells, the blebs observed in this study did not retract and were stable during an imaging time of 90 min (Fig. S5). The nearly linear increase of Sytox Green staining of nuclei upon MbxA treatment was steeper if higher toxin concentrations were applied (Fig. 10). This indicates that the cell membrane is immediately permeabilized after contact with MbxA and possibly the number of pores increases with increasing toxin concentrations. This is also in clear contrast to RtxA1, which displays a cytotoxicity that depends on contact between the bacterium and the host cell (59). Additionally, accumulation of labeled Atto488-MbxA^S9C^ on HEp-2 cell membranes is an indication of the presence of potential receptors for MbxA binding on human epithelial cells, which might be involved in pore formation in HEp-2 cells. The CD18 subunit of the ß2 integrin was found to be the receptor for RTX toxins such as *Actinobacillus actinomycetemcomitans* leukotoxin and *E. coli* HlyA, promoting cytotoxicity against human immune cells (34, 35). Presence of receptors on HEp-2 cells (87), together with our observation of accumulation of Atto488-MbxA^S9C^ on HEp-2 cell membranes further enhances the possibility of a potential receptor for MbxA on HEp-2 cells as well.

In summary, we demonstrated that the *M. bovis* cytotoxin MbxA is secreted and acylated by the heterologous UPEC HlyA system in an *E. coli* lab strain, which allowed purification of proMbxA and MbxA from *E. coli* culture supernatants. *E. coli* UTI89 HlyC efficiently acylated

MbxA at the two predicted lysine residues with predominantly C_14_ fatty acids. The acylation of MbxA is strictly necessary for activity against sheep RBC, human epithelial and human T cells, as proMbxA did not exert any cytotoxicity. The cytotoxicity studies showed that MbxA is not species- or cell type-specific and active against human cells in a nanomolar range. Live-cell imaging revealed localization of MbxA at the plasma membrane of human epithelial cells, an immediate permeabilization and membrane blebbing upon contact with MbxA.

## Material and Methods

### Sequence alignment and topology prediction

Alignment was carried out using EMBOSS Needle pairwise sequence alignment (88). For the prediction of membrane spanning regions the Constrained Consensus TOPology prediction server (CCTOP) with different implemented algorithms was used (52).

### Cloning of the *mbxA* gene

The *mbxA* gene (Uniprot Q93GI2) was amplified from *M. bovis* DSM 6328 genomic DNA with the primer pair mbxA-for 5‘-AACCTTTTCTAACACAACGAGGAGAGAC-3‘ and mbxA-rev 5‘- AAATCACTAAACACTTGGAGCCAAAATTC-3‘ and cloned into the pJET1.2 vector (Thermo Scientific). Subsequently the *mbxA* gene was cloned into the pSU2726 *hlyA* vector (60) replacing the *hlyA* gene. An N-terminal His_6_-Tag site was introduced, resulting in plasmid pSU- *6H-mbxA*. To produce acylated MbxA, *6H-mbxA* was cloned into the pSU-*hlyC-6H*/*hlyA* plasmid (67) replacing the *hlyA* gene. To restore the *hlyA* enhancer region (89) the His_6_-Tag of *hlyC* was deleted, creating pSU-*hlyC/6H-mbxA.* In this vector, both genes, *hlyC* and *mbxA*, form a bicistronic operon and are under the control of one LacZ promoter (67).

### Expression and secretion of proMbxA and MbxA

Secretion of MbxA in *E. coli* BL21(DE3) was facilitated by co-expression of the transporter components HlyB and HlyD of the *E. coli* UIT89 HlyA system. For expression of recombinant proMbxA, ten 300 ml flasks containing 50 ml LB with 100 µg/ml ampicillin and 30 µg/ml kanamycin per flask were inoculated with an *E. coli* BL21(DE3) overnight culture harboring plasmids pK184-*hlyBD* (13) and pSU 6H-*mbxA* to obtain a starting OD_600_ of 0.05. The culture was grown at 37°C and 180 rpm until OD_600_ reached 0.4-0.6. Subsequently, the culture was supplemented with 10 mM CaCl_2_ and expression was induced with 1 mM IPTG. After 5 h of expression at 37°C and 180 rpm the cells were removed by centrifugation at 13500 g for 45 min. The supernatant was filtered with a 0.45 µm filter and stored on ice for purification. For the secretion of active MbxA, *E. coli* BL21(DE3) transformed with the plasmids pK184- *hlyBD* and pSU-*hlyC/6H-mbxA* were grown in 2 L LB medium with 100 µg/ml ampicillin and 30 µg/ml kanamycin in baffled 5 L flasks at 37°C and 160 rpm and induced at the same conditions as pSU-*6H-mbxA*. Secreted MbxA accumulated in the foam of the culture. After the culture reached an OD_600_ between 3.5-4.0, the foam was collected and centrifuged at 4000 g for 10 min. The condensed foam was solubilized in solubilization buffer (6 M urea, 100 mM NaCl, 50 mM TRIS pH 8.0) by stirring at room temperature overnight. Aggregate was collected by centrifugation at 160.000 g for 30 min and the solubilized protein stored at -80°C.

### Purification of MbxA

Solubilized MbxA was refolded by dropwise dilution from 6 M to 400 mM urea with IMAC buffer (50 mM Tris pH 7.8, 400 mM NaCl, 10 mM CaCl_2_) with constant stirring at room temperature. All subsequent steps were performed at 4°C. Refolded MbxA was loaded on a 5 ml Ni^2+^ loaded HiTrap IMAC HP column (GE Healthcare) and eluted with a linear 0-75 mM histidine gradient with elution buffer (50 mM Tris pH 7.8, 400 mM NaCl, 10 mM CaCl_2,_ 75 mM histidine). Elution fractions containing MbxA were pooled and concentrated with Amicon Ultra-15 Centrifugal Filter Units (50 kDa NMWL, Merck Millipore). Histidine was removed with a PD10 desalting column (GE Healthcare) at room temperature and the protein eluted with cold IMAC buffer. Activity of MbxA was tested on Columbia agar with 5% sheep blood (Oxoid).

### Purification of proMbxA

The culture supernatant containing proMbxA was concentrated with Amicon Ultra-15 Centrifugal Filter Units (50 kDa NMWL, Merck Millipore) to 1/10 of the starting volume and loaded on a 5 ml Ni^2+^ HiTrap IMAC HP column (GE Healthcare). A linear 0-75 mM histidine gradient with elution buffer was used to elute proMbxA. Elution fractions containing proMbxA were pooled and concentrated. Histidine was removed with a PD10 desalting column (GE Healthcare) and the protein eluted with IMAC buffer.

### Size Exclusion Chromatography (SEC) and Multi Angle Light Scattering (MALS)

For SEC, proMbxA or MbxA containing IMAC fractions were pooled, concentrated to 0.5 ml and after centrifugation at 21,700 g for 20 min directly applied to a Superose 6 Increase 10/300 GL column (GE Healthcare) in SEC buffer (50 mM TRIS pH 7.8, 100 mM NaCl, 10 mM CaCl_2_). The chromatography was performed on an ÄKTA Purifier system (GE Healthcare). proMbxA peaks resulting from the SEC were pooled separately and stored with 20% glycerol at -80°C. A subsequent SEC-MALS analysis was carried out to investigate the oligomeric state of the proMbxA species. The separated species were concentrated to 1.3 - 2.5 mg/ml and 200 µl of each were applied to a Superose 6 Increase 10/300 GL column (GE Healthcare) equilibrated with SEC buffer and connected to a miniDAWN TREOS II triple-angle light scattering detector (Wyatt Technologies) and an Optilab T-rEX differential refractive index detector (Wyatt Technologies). The set up was performed using an Agilent 1260 HPLC system with a flowrate of 0.6 ml/min and the data was analyzed with the ASTRA 7.1.2.5 software (Wyatt Technologies).

### Expression and purification of proHlyA and HlyA

proHlyA was expressed in *E. coli* BL21(DE3) carrying pSU2726-*hlyA* (60) without transporter components to produce inclusion bodies. 2 L of 2xYT medium with 100 µg/ml ampicillin in baffled 5 L flasks were inoculated with an overnight culture to OD_600_ 0.1 and incubated at 37°C and 160 rpm. At an OD_600_ of 0.6 expression was induced with 1 mM IPTG, and expression continued for 4 h at 37°C and 160 rpm. Afterwards cells were collected by centrifugation for 15 min at 13900 g and inclusion bodies purified according to (90). proHlyA solubilized in 6 M urea was refolded by dropwise dilution to 400 mM urea with 100 mM HEPES pH 8.0, 250 mM NaCl, 10 mM CaCl_2_. The refolded protein was concentrated with Amicon Ultra-15 Centrifugal Filter Units (100 kDa NMWL, Merck Millipore) and remaining urea removed with a PD10 desalting column (GE Healthcare). The eluted protein was concentrated and applied to a Superose 6 Increase 10/300 GL column (GE Healthcare) in the same buffer.

Active HlyA was purified from secreting *E. coli* BL21(DE3) harboring pK184-*hlyBD* and pSU-6H- *hlyC/hlyA*. 2 L 2xYT medium with 100 µg/ml ampicillin and 30 µg/ml kanamycin in baffled 5 L flasks were inoculated with an overnight culture to OD_600_ 0.1 and grown at 37°C and 160 rpm. At OD_600_ 0.4-0.6 the culture was supplemented with 10 mM CaCl_2,_ and expression induced with 1 mM IPTG. After 4 h of expression the foam of the culture was collected, HlyA solubilized in 6 M urea and refolded by dropwise dilution as described for MbxA. Refolded HlyA was concentrated (Amicon Ultra-15 Centrifugal Filter Units, 100 kDa NMWL, Merck Millipore) and remaining urea removed with PD10 desalting column (GE Healthcare).

### MS analysis

MbxA, proMbxA as well as HlyA and proHlyA were separated in a polyacrylamide gel, stained with Coomassie Brilliant Blue (CBB), the corresponding bands of the proteins were cut out from the gel and processed for mass spectrometric analysis as described (91). Briefly, proteins were reduced with dithiothreitol, alkylated with iodoacetamide, and digested with trypsin. Resulting peptides were separated during a 1 h gradient on C18 material using an Ultimate3000 rapid separation liquid chromatography system (Thermo Fisher Scientific) and subsequently injected in an online-coupled QExactive plus mass spectrometer (Thermo Fisher Scientific) via a nano-electrospray interface (described in (91)). Briefly, survey scans were recorded at a resolution of 70,000 and subsequently up to 10 precursors were selected by the build-in quadrupole, fragmented via higher energy collisional dissociation, and analyzed at a resolution of 17,500.

Recorded spectra were further analyzed by MaxQuant version 1.6.10.43 (MPI for Biochemistry, Planegg, Germany) using standard parameters if not stated otherwise. Searches for MbxA and proMbxA were carried out in a dataset containing 2734 *Moraxella bovis* sequences (UP000254133, downloaded on 27_th_ February from the UniProt KB) as well as an entry for MbxA. For HlyA and proHlyA, a data set was used containing 4156 *Escherichia coli* BL21(DE3) sequences (UP000002032, downloaded on 29^th^ April 2019 from the UniProt KB)and an additional entry for HlyA. Carbamidomethylation at cysteines was considered as fixed and methionine oxidation, n-terminal acetylation, and acylation with C_12_, C_13_, C_14_, C_15_, C_16_, C_17_ and C_18_ chains at lysines as variable modifications. In a second search, additionally the hydroxylated variants of the acyl chains were considered. The minimal peptide length was set to five, and two variable modifications were allowed per peptide. Identified proteins and peptides were reported at a false discovery rate of 1%.

### Cell culture

Human epithelial cells, HEp-2 (epithelial larynx carcinoma, ATCC-CCL-23) were routinely cultured in Dulbeccós modified Eaglés medium (DMEM GlutaMAX™; Thermo Fisher) supplemented with 10% fetal calf serum (FCS), MEM vitamins, non-essential amino acids, amphotericin B (2.5 µg/ml) and gentamicin (50 µg/ml) (all Thermo Fisher Scientific) and human T lymphocytes, Jurkat J16 (Adult lymphoblastic leukemia ACC-282, DSMZ) were cultured in RPMI-1640 (Thermo Fisher) supplemented with 10% FCS, 100 U/ml penicillin, 100 µg/ml streptomycin and 10 mM HEPES at 37°C under 5% CO_2_.

### Lactate dehydrogenase (LDH) release assay

To monitor the cytotoxicity of MbxA the release of cytosolic LDH was measured as an indicator of membrane damage using the LDH Cytox Assay Kit (Biolegend).

For HEp-2 cells, 10,000 cells per well were seeded in a 96 well plate and grown over night. Prior to the assay, the medium was exchanged to DMEM supplemented with 5% FCS. For Jurkat cells, 100 µl with a density of 6×10^5^ cells/ml in RPMI with 5% FCS were seeded in 96 well plates and for both cell lines all following incubations were carried out in the appropriate cell culture medium supplemented with 5% FCS. HEp-2 cells and Jurkat cells were incubated with a serial dilution of MbxA in DMEM respectively RPMI for 1 h at 37°C under 5% CO_2_. To assess if preincubation with proMbxA influences MbxA cytotoxicity purified proMbxA was added to the HEp-2 cells together with fresh DMEM and incubated for 30 min at 37°C under 5% CO_2_ before addition of the serial dilution of MbxA. For Jurkat cells proMbxA was added to the cell suspension immediately before seeding and the cells were likewise incubated for 30 min at 37°C under 5% CO_2_ before the addition of MbxA. To monitor the basal release and activity of LDH, HEp-2 and Jurkat cells were incubated with cell culture medium with IMAC buffer (basal LDH). Plates were centrifuged (3 min, 1000 g) and supernatants of HEp-2 and Jurkat cells were collected. For the quantification of the LDH release, the assay was carried out according to manufactureŕs manual (Biolegend) and absorbance measured at 490 nm. The assay was conducted in biological and technical triplicates. For the calculation of the cytotoxicity absorbance of the background control was subtracted from all other values and the average absorption of triplicates was calculated. The relative cytotoxicity was normalized to the maximal LDH release (maximal absorption after MbxA treatment, Max Abs (MbxA treated)) of each experiment and calculated with the following equation:

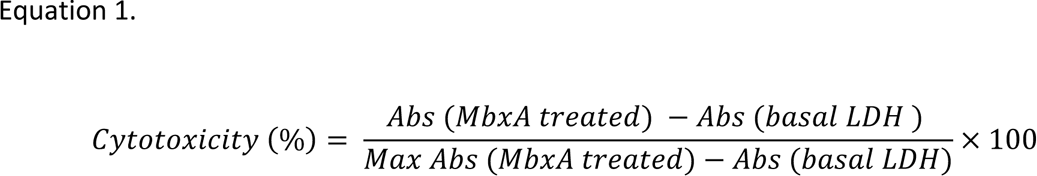

The concentration of the half-maximal effect of MbxA (cytotoxic dose 50, CD_50_) was calculated from three biological replicates with GraphPad Prism 7 with the following equation: Equation 2.

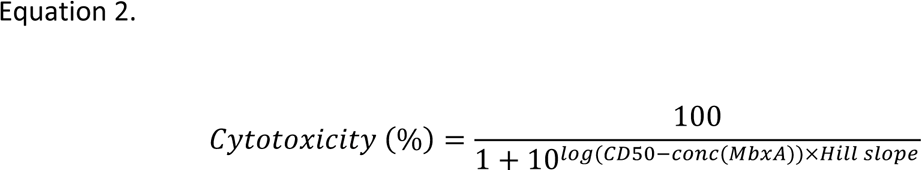

### Labelling of MbxA with Atto-488 maleimide

Since MbxA does not have any intrinsic cysteine, a serine to cysteine point mutation (StoC) in the N-terminal region of MbxA at the 9^th^ amino acid position was introduced. Site directed mutagenesis of the plasmid pSU-hlyC/6H-mbxA was performed by PCR using the primer pairs mbxAStoC-for 5’-GTAATTAAAT**<u>G</u>**TAATATTCAAGCAGG-3’ and mbxAStoC-rev 5’- ATTTATATTGGACATACGAC-3’. Serine to cysteine point mutation was confirmed by sequencing and the lytic activity of the mutated MbxA was tested on Columbia agar with 5% sheep blood (Oxoid). For the quantitative comparison of MbxA activity, hemoglobin release assay was performed (protocol given below).

Secretion and purification of MbxA^S9C^ was performed using the protocol for wild type MbxA as described above. Labeling reaction with Atto-488 was performed based on manufacturer’s instructions for thiol-reactive fluorescent dyes. 6 ml of purified MbxA^S9C^ with a concentration of 0.5 mg/ml was used for labeling reactions. MbxA^S9C^ was incubated with 10-fold molar excess of TCEP (0.5 M TCEP stock solution) for 6 hours on ice. The solution was concentrated to 2.5 ml using a 50 kDa MWCO filter concentrator, buffer was exchanged to DPBS buffer (Solution C – see below) using a PD10 column and the solution was again concentrated to 4 mg/ml (540 µL) using a 50 kDa MWCO filter concentrator. Subsequently, reduced MbxA^S9C^ was incubated with a 15-fold molar excess of Atto-488 (5 mM stock solution in DMSO) for 24 hours on ice. The reaction was quenched using a 5-fold molar excess of GSH (6 mM stock solution in DPBS++ buffer (Solution A – see below)) compared to the dye and incubated for 10 minutes on ice. Unbound dye and GSH were removed using a PD10 column buffered with DPBS++ buffer (Solution A – see below). Furthermore, the in-gel fluorescence of the labeled protein was detected by a Bio-Rad gel documentation system (Fig.S3).

[DPBS++ buffer (**Solution A**): 0.90 mM CaCl_2_.2H_2_O, 0.5 mM MgCl_2_.6H_2_O, 2.67 mM KCl, 1.47 mM KH_2_PO_4_, 137.9 mM NaCl, 8.06 mM Na_2_HPO4, pH 7.4. **Solution B**: 0.2 M sodium bicarbonate solution, adjusted to pH 9.0 with 2 M sodium hydroxide. Labeling buffer (**Solution C**) --> To 20 parts of Solution A add 1 part of Solution B to obtain a labeling buffer of pH 8.3.

### Calculation of Degree of Labeling (DOL)

The degree of labeling (DOL, dye-to-protein ratio) was determined by absorption spectroscopy (Table S2: Absorbance (A) = extinction coefficient (ε) × molar concentration (c) × path length (d). Labeling efficiency/ DOL = Concentration of the dye/ Concentration of the protein.

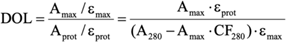

Absorbance of the labeled dye at 494 nm and 280 nm were measured using a NanoDrop^TM^. As all dyes show some absorption at 280 nm, the measured absorbance A_280_ must be corrected for the contribution of the dye. This is given by Amax × CF_280_.

### Hemoglobin Release Assay

Defibrinated sheep blood was obtained from Oxoid. 1 ml of cells were transferred into 1.5 ml reaction tubes and centrifuged for 1 min at 14,000 g. The supernatant was removed, and cells were resuspended in buffer containing 10 mM Tris-HCl pH 7.5, 155 mM NaCl, 20 mM CaCl_2_, 5 mM KCl and 2 mM MgSO_4_ by 5-10 sec vortexing intervals. This process was repeated until the supernatant remained colorless. The resuspended cells were transferred into a 50 ml falcon tube to guarantee equal cell distribution and immediately used for the *in vitro* assay. All steps were performed at room temperature. 250 nM concentrations of Atto-488 labeled MbxA^S9C^, MbxA^S9C^, MbxA or HlyA were added to 500 μl washed sheep blood cells each. 16% SDS was used as the positive control and set arbitrarily as 100% and IMAC buffer was used as the negative control. The cells were incubated for 30 minutes at 37°C with the proteins. Subsequently, the cells were removed by centrifugation (1 min at 14,000 x g) and 200 μl of 1/10 diluted fraction of the supernatant transferred to a 96-well plate. The hemoglobin amount in the supernatant was determined on a FLUOstar OPTIMA microplate reader (BMG Labtech) with an absorption wavelength of 544 nm.

### Live-cell imaging

HEp-2 cells were cultivated and passaged in DMEM (Pan Biotech) supplemented with 10% FCS, MEM vitamins, non-essential amino acids, amphotericin B (2.5 µg/ml) and gentamicin (50 µg/ml) (all Thermo Fisher Scientific). Two days prior to use, HEp-2 cells were seeded in 2 ml DMEM in 35 mm μ-Dish 1,5H glass bottom dishes (Ibidi) and grown for 39 - 48 h at 37 °C under 5 % CO_2_.

In case of the incubation experiment with unlabeled MbxA, cells were washed twice with 1 ml of prewarmed DMEM supplemented with 20 mM HEPES as live cell medium (Pan Biotech) and preincubated with 250 µl of 20 µM of Hoechst 33342 (Invitrogen) in live cell medium for 22 min at 37 °C and 5 % CO_2_. After this, 750 µl of live cell medium containing 1 µl of 5 mg/ml Hoechst 33342 (Invitrogen), resulting in a 5 µM concentration of Hoechst 33342 and 5 µg/ml CellMask™ Deep Red and were incubated for another 8 min at 37 °C and 5 % CO_2_ (total incubation of 30 min). The cells were washed once gently with 1 ml of pre-warmed live cell medium and 1 ml of live cell medium with a 5 µM concentration of Sytox® Green was added. The cells were transferred to a Zeiss LSM880 Airyscan microscope system (Carl Zeiss Microscopy GmbH). A region of interest was preset for the incubation experiment. 1 ml of a fresh solution of MbxA or proMbxA protein was prepared at the 2-fold of the indicated final concentration in 1 ml of live cell medium containing a 5 µM concentration of Sytox Green. The MbxA solution at twice the concentration was added to the cells and gently mixed by pipetting three times without touching the live cell dish to keep the region of interest in focus. This resulted in total in 2 ml live cell medium containing the indicated final concentration of MbxA supplemented with 5 µg/ml of Sytox® Green during the acquisition of the time series. The measurement was started immediately.

In the case of incubation experiment with Atto488-labeled MbxA^S9C^, cells were washed with 1 ml of DMEM supplemented with 20 mM HEPES as live cell medium (Pan Biotech, Germany), incubated in 1ml of pre-warmed live cell medium and transferred to the LSM880 (Carl Zeiss Microscopy GmbH) focusing a region of interest. 1ml of a fresh solution of Atto488-labeled MbxA^S9C^ at twice the final concentration was prepared in live cell medium, added and again gently mixed by pipetting three times without touching the live cell dish to keep the region of interest in focus. The measurement was started immediately. After 30 min the supernatant containing the Atto488-labeled MbxA^S9C^ was carefully replaced by fresh pre-warmed live cell medium and analyzed for protein localization again.

The imaging conditions were set up as follows. Confocal and Airyscan micrographs were recorded using a Zeiss LSM880 Airyscan microscope system (Carl Zeiss Microscopy GmbH) equipped with a Plan-Apochromat 63x/1.4 oil immersion objective lens. For excitation a 405 nm Laser was used for Hoechst 33342, a 488 nm Argon laser for excitation of Sytox Green and Atto488-labeled MbxA^S9C^ and a 633 nm laser for excitation of CellMask Deep Red.

In confocal microscopy of unlabeled MbxA a pixel dwell time of 1 µs was used without averaging at a frame rate of 15 sec/frame to reduce phototoxicity to a minimum. Excitation and detection of different dyes was set up in a frame wise switch to reduce crosstalk and detection ranges were set for Hoechst 33342 at 415 – 460 nm, for Sytox Green at 495 – 550 nm and for CellMask Deep Red at 648 - 700 nm. Focus was maintained during the time series using the hardware based autofocus system Definite Focus 2 (Carl Zeiss Microscopy GmbH).

In confocal microscopy of Atto488-labeled MbxA^S9C^ tiles can experiments of 25 single images were acquired at a pixel dwell time of 0.77 µs and the detection range for ^Atto488^MbxA was set at 495 – 550 nm. Additional super-resolution Airyscan micrographs were acquired at a pixel dwell time of 2 µs and a 4-fold averaging in Airyscan super-resolution mode after alignment on detector elements was established. To limit the detection range a BP 495-535 + LP 555 filter was used. The final Airyscan image was generated by using the Zeiss Airyscan processing at 2D automatic settings.

### Image Quantification

Confocal images of unlabeled MbxA were analyzed using Cellprofiler image analysis software version 4 (92). The nuclei area were determined by merging the Hoechst 33342 and Sytox Green signal and applying adaptive Sauvola thresholding. Tracking of nuclei across the time series was performed by the TrackObjects module with overlap tracking and a maximum pixel distance of 40 to consider matches. Extracted image data were processed to consider only nuclei that have been successfully tracked during the whole time series of 20 minutes (80 frames). The Cellprofiler pipeline and Jupyter notebook for data processing are available publicly at github (https://github.com/abhamacher/timelapseMic_MbxA).

## Acknowledgments

We thank Diana Kleinschrodt for valuable support and all members of the Institute of Biochemistry for fruitful discussions. We would also like to thank former students Nicole Jasny and Fabian Adamek. This study was funded by the Jürgen Manchot Graduate School “Molecules of Infection III & IV” (to J.H. and L.S.), by the RTG 2158 (to S.W.) and the RTG 2578 (to S.W.) and CRC 1208 (project INF to S.W.-P.) of the Deutsche Forschungsgemeinschaft.

## Conflict of interest

The authors declare no conflict of interest.

## Data availability

All fluorescence data have been deposited at BioImage Archive (https://www.ebi.ac.uk/biostudies/studies/S-BIAD295). The mass spectrometry proteomics data have been deposited to the ProteomeXchange Consortium via the PRIDE (93) partner repository with the dataset identifier PXD030929. All other data are available upon request by the authors.

## Notes

### Competing Interest Statement

The authors have declared no competing interest.

