## Supplementary Information for "Heterologously secreted MbxA from *Moraxella bovis* induces a membrane blebbing response of the human host cell"

```
HlyA 544 KFVTPLLTPGEEIRERRQSGKYEYITELLVKGVDKWTVKGV 584
         .|.:||||.|.|.|||..:|||.||.:|....|..|.|..
MbxA 516 HFTSPLLTAGTESRERLTNGKYSYINKLKFGRVKNWQVTD- 555

HlyA 670 LGGDVKVLQEVVKEQEVSVGKRTEKTQYRSYEFTHINGKNL 710
         ..||  :..||||.||..||||||..|||.||...: |...
MbxA 642 ARGD--IYHEVVKRQETKVGKRTETIQYRDYELRKV-GYGY 679
```

**Fig. S1** Sequence alignment (1) of the acylation sites of HlyA sites at lysine residues K564 and K690, which are highlighted in bold (2). The homologous residues K536 and K660 predicted to be acylated in MbxA (3) are likewise shown in bold.

**Table S1:** MS analysis of MbxA acylation. MbxA was *in vivo* cross-acylated by co-expressing HlyC (4). Modified peptides are listed with their detected fatty acid and hydroxy fatty acid modifications ($C_{12}$-$C_{16}$) and the corresponding number of peptide spectrum matches. Mass shifts contributing to hydroxy fatty acids could possibly originate from the oxidation of tyrosine residues in the peptide fragment and PSM of supposed hydroxy acylations are therefore marked (*). The acylated lysine residues are shown in bold in row 'Sequence'. Furthermore, the charge states are given with which the respective peptides were detected in the mass spectrometer as well as the score from search engine MaxQuant.

Acyl modifications were detected only in peptides covering the predicted lysine residues K536 and K660, respectively. Only a low number of PSM were detected for K536 and no K660 lysine containing peptides in proMbxA. This is probably due to a higher specific cleavage of the respective peptides resulting in quite small fragments which have not been considered in the search. Here, the cleavage behind K536 and K660 might not be masked by acylation of the respective residues. The analysis of the *in vitro* acylation of HlyA by HlyC confirms the acylation of the K564 and K690 acylation sites.

| Sequence | Site | Modifications | Charges | Score | PSM |
|---|---|---|---|---|---|
| **MbxA** | | | | | |
| ERLTNGKYSYINK | K536 | C14 | 2 | 209.19 | 5 |
| ERLTNGKYSYINK | K536 | C14 - OH* | 2 | 103.22 | 2 |
| ERLTNGKYSYINK | K536 | C15 | 3 | 61.423 | 1 |
| LTNGKYSYINK | K536 | C12 -OH* | 2 | 50.909 | 1 |
| LTNGKYSYINK | K536 | C13 | 2 | 93.649 | 2 |
| LTNGKYSYINK | K536 | C14 | 2 | 150.66 | 17 |
| LTNGKYSYINK | K536 | C14 - OH* | 2 | 63.159 | 1 |
| LTNGKYSYINK | K536 | C15 | 2 | 139.98 | 4 |
| LTNGKYSYINK | K536 | C16 | 2 | 56.258 | 1 |
| LTNGKYSYINKLK | K536 | C14 | 2 | 146.11 | 2 |
| LTNGKYSYINKLK | K536 | C14 - OH* | 2;3 | 69.815 | 2 |
| VGKRTETIQYR | K660 | C14 | 3 | 69.721 | 1 |
| **proMbxA** | | | | | |
| LTNGKYSYINKLK | K536 | Unmodified | 2;3;4 | 299.29 | 6 |
| **HlyA** | | | | | |
| QSGKYEYITELLVK | K564 | C14 | 2 | 92.19 | 1 |
| QSGKYEYITELLVK | K564 | C14 - OH* | 2 | 74.611 | 1 |
| QSGKYEYITELLVK | K564 | Unmodified | 3 | 83.633 | 4 |
| RQSGKYEYITELLVK | K564 | C14 | 3 | 90.754 | 1 |
| RQSGKYEYITELLVK | K564 | C14 - OH* | 3 | 93.909 | 1 |
| RQSGKYEYITELLVK | K564 | Unmodified | 3 | 71.545 | 1 |
| EQEVSVGKR | K690 | C12 | 2 | 49.813 | 1 |
| EQEVSVGKR | K690 | C14 | 2;3 | 190.08 | 3 |
| EQEVSVGKR | K690 | C14 - OH* | 2 | 159.04 | 3 |
| VLQEVVKEQEVSVGKR | K690 | C13 | 3 | 76.85 | 2 |
| VLQEVVKEQEVSVGKR | K690 | C14 | 2;3 | 314.04 | 45 |
| VLQEVVKEQEVSVGKR | K690 | C14 - OH* | 2;3 | 173.11 | 5 |
| VLQEVVKEQEVSVGKR | K690 | C16 | 3 | 97.836 | 1 |
| VLQEVVKEQEVSVGKR | K690 | Unmodified | 2 | 75.479 | 2 |
| VLGGDVKVLQEVVKEQEVSVGK | K690 | Unmodified | 3 | 19.975 | 1 |
| VLQEVVKEQEVSVGK | K690 | Unmodified | 2;3 | 126.83 | 4 |
| **proHlyA** | | | | | |

| | | | | | |
|---|---|---|---|---|---|
| QSGKYEYITELLVK | K564 | Unmodified | 2;3 | 287.76 | 36 |
| RQSGKYEYITELLVK | K564 | Unmodified | 2;3 | 158.11 | 32 |
| EQEVSVGKRTEK | K690 | Unmodified | 2;3 | 122.51 | 4 |
| VLGGDVKVLQEVVKEQEVSVGK | K690 | Unmodified | 2;3;4 | 241.13 | 5 |
| VLQEVVKEQEVSVGK | K690 | Unmodified | 2;3 | 283.47 | 69 |
| VLQEVVKEQEVSVGKR | K690 | Unmodified | 2;3;4 | 334.04 | 36 |

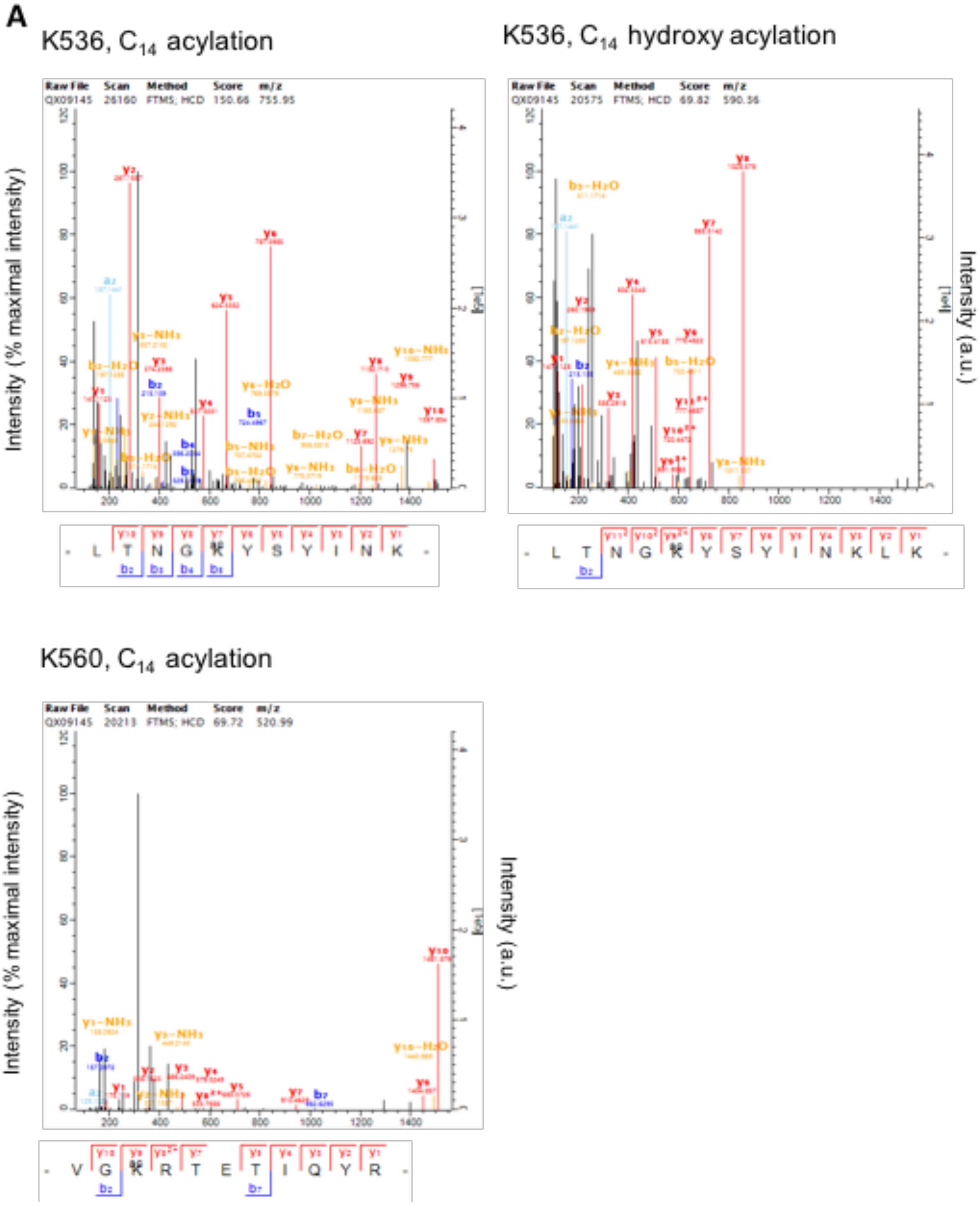
A
K536, C14 acylation
K536, C14 hydroxy acylation
K560, C14 acylation
Raw File Scan Method Score m/z
QX09145 26160 FTMS; HCD 150.66 755.95
QX09145 20575 FTMS; HCD 69.82 590.56
QX09145 20213 FTMS; HCD 69.72 520.99
Intensity (% maximal intensity)
Intensity (a.u.)
L T N G K Y S Y I N K
L T N G K Y S Y I N K L K
V G K R T E T I Q Y R

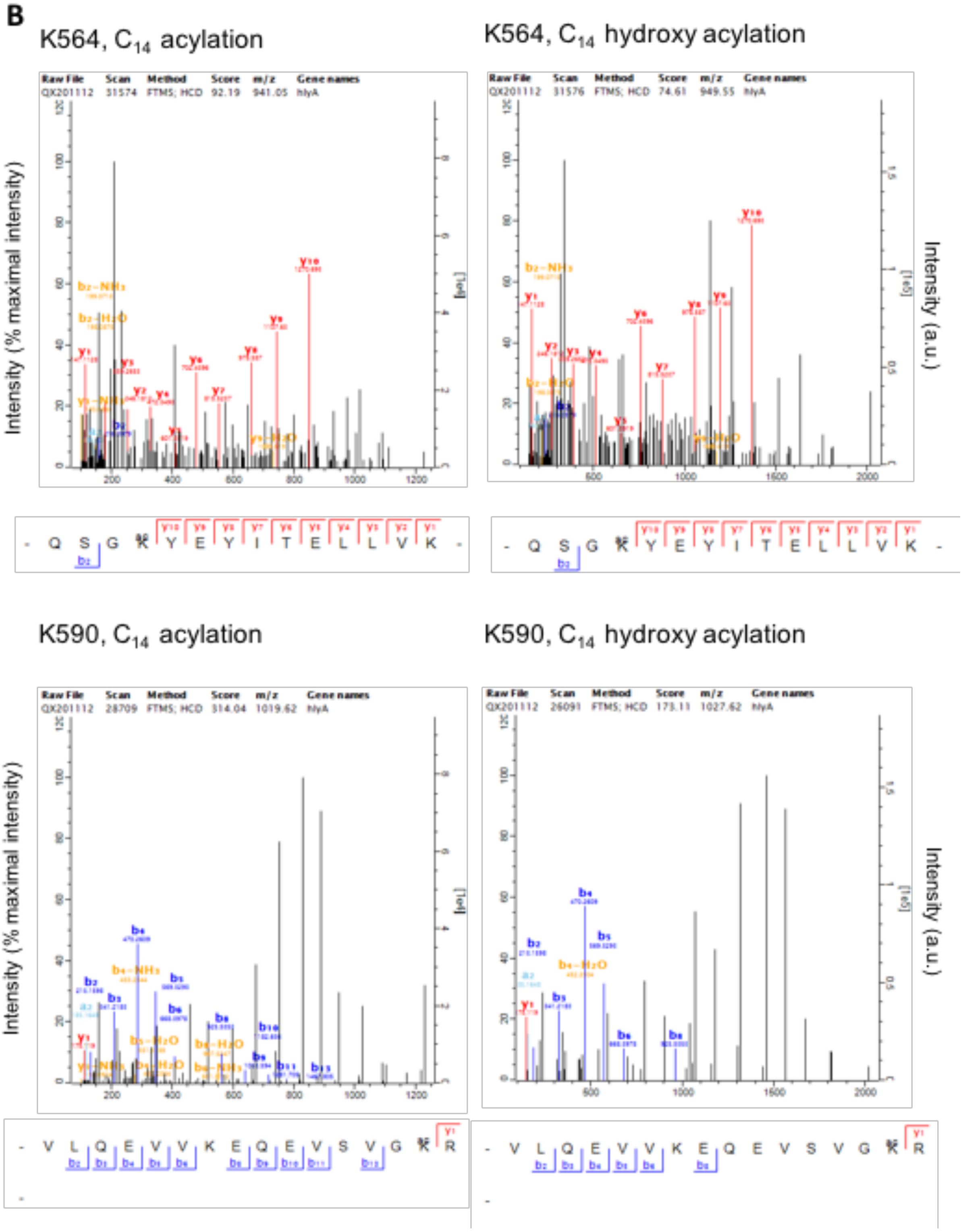


**Fig. S2 Exemplary** MS spectra of the peptides that cover the first, K536, and second acylation site, K660, of MbxA (A) and of the two acylation sites K564 and K690 of HlyA (B) modified with $C_{14}$ or $C_{14}$-OH* acylation.

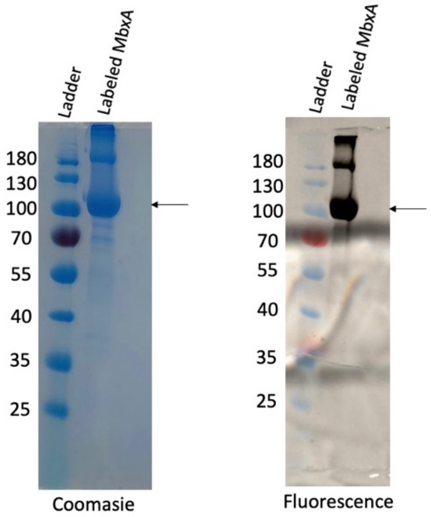


**Fig. S3**. Coomassie staining and Fluorescence of $^{Atto488}$MbxA on SDS gel. Labelling of MbxA with Atto-488 was confirmed on SDS PAGE by Coomassie staining and Fluorescence. The fluorescence of the $^{Atto488}$MbxA was detected after separation in the SDS gel by a BioRad gel documentation system. Arrow mark shows the $^{Atto488}$MbxA monomers. The protein dimers near 180 kDa are more prominent due to the introduced Cysteine and subsequent disulfide bond formation.

| **Parameters** | **Values** |
|---|---|
| Molecular weight of MbxA (Da) | 98,846 |
| Extinction coefficient of MbxA at 280 nm in $H_2O$ ($M^{-1}cm^{-1}$) | 58000 |
| Absorbance value at 280 nm | 1.34 |
| Absorbance value at 494 nm (Atto-488) | 2.3 |
| Specific correction factor for the Atto-488 | 0.09 |
| **Protein concentration (mg/ml)** | 0.46 (3.4 mL) |
| Extinction coefficient of Atto-488 at 494 nm ($M^{-1}cm^{-1}$) | 90000 |
| **Labeling efficiency (fraction)** | 1.308 |

**Table S2:** Calculation of Labeling efficiency of MbxA with Atto-488 maleimide. Absorbance values at 280 nm and 494 nm are measured using NanoDrop™ One. The evaluation gives a protein concentration of 0.46 mg/ml and a labeling efficiency of 130.8% for the protein labeled with Atto-488 maleimide.

Using epifluorescence / DIC microscopy, experiments were performed using another set of dyes. To visualize the changes in the membrane morphology, the HEp-2 plasma membrane was labeled with Wheat Germ Agglutinin conjugated with Alexa Fluor 488 (WGA-488) and to monitor possible permeabilization of the cells, propidium iodide (PI) fluorescence was measured. As observed in confocal microscopy, membrane blebbing and permeabilization of HEp-2 cells observed in epifluorescence / DIC microscopy as well, when treated with MbxA, but not with proMbxA (Fig. S4). Incubation of HEp-2 cells with proMbxA did not show any visible changes in the membrane morphology over a period of 20 min. Additionally, no increase in PI signal intensity was detected indicating preserved plasma membrane integrity. HEp-2 cells were incubated with different concentrations of MbxA (250 nM, 30 nM and 10 nM) and proMbxA (250 nM), and images were taken in every 10 s over a total time period of 20 min (Fig. S4). Incubation of HEp-2 cells with 250 nM MbxA resulted in membrane bulges in first few minutes itself. At the same time, PI influx and DNA staining also started occurring, and the staining increased with higher incubation time of the same experiment (Fig. S5). As observed in confocal microscopy, 30 nM MbxA incubation of HEp-2 cells also resulted in the membrane blebbing and PI influx, but delayed and showing lower intensity response when compared to 250nM MbxA treatment (Fig. S4 and Fig. S5). In the case of 10 nM MbxA, smaller membrane bulges were visible only after 20 min going in hand with a small slope of PI fluorescence signal increase. After an extended incubation time of 90 min, small bulges as well as large blebs of approximately 10 µm were visible and PI staining could be detected demonstrating the dose dependent effect of cell permeabilization by MbxA again.

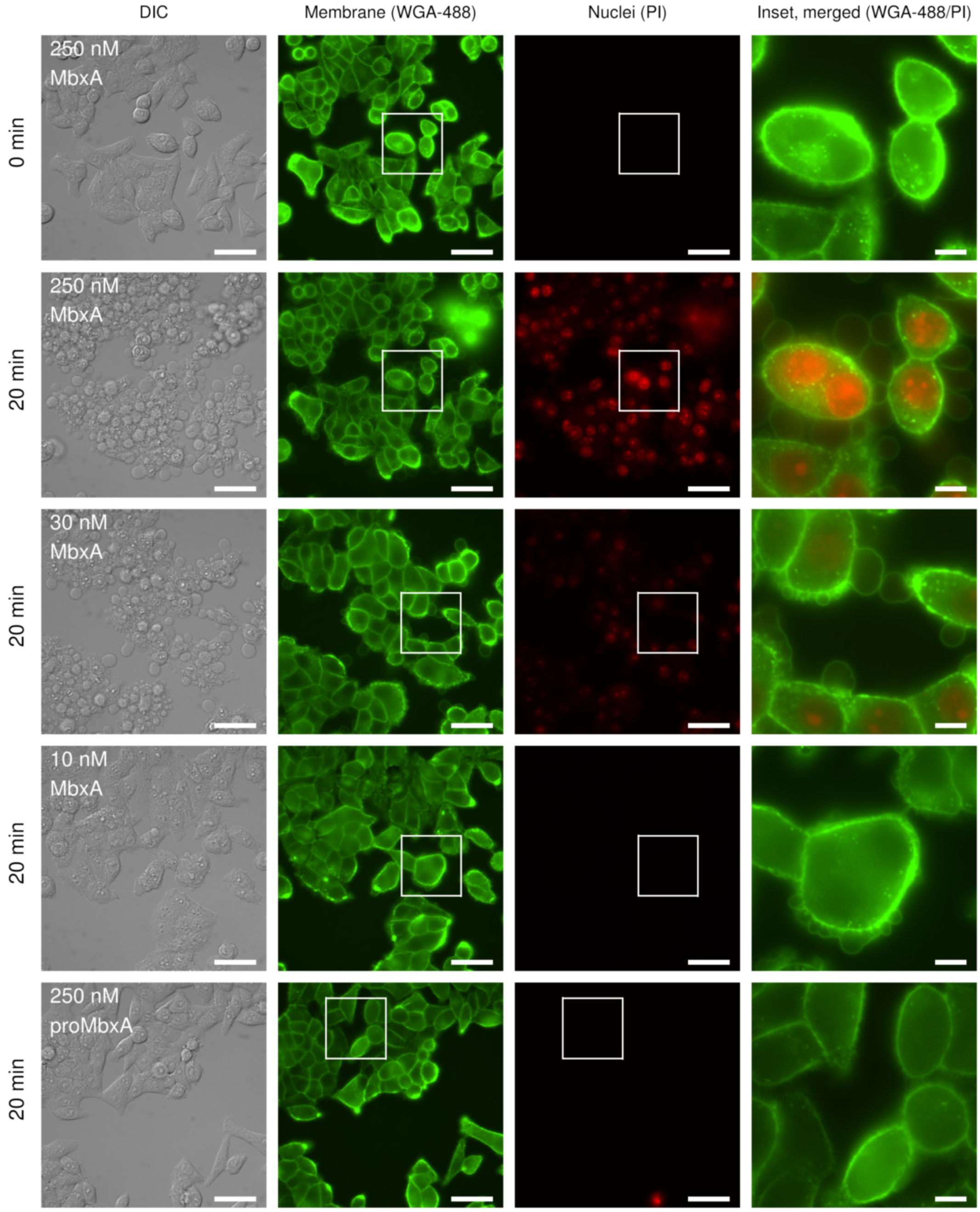


**Fig. S4**. Epifluorescence live-cell imaging of HEp-2 cells treated with 250 nM, 30 nM or 10 nM of MbxA and 250 nM of proMbxA for 20 min. HEp-2 cells exposed to 250 nM of MbxA are shown after 0 min and after 20 min of incubation (first and second row). HEp-2 cells treated with 30 nM and 10 nM are shown after 20 min (third and fourth row). Data at later timepoints of the same experiment confirm potential permeabilization at lower concentrations in Fig.S5. HEp-2 cells incubated with 250 nM proMbxA and images after 20 min (fifth row). Membranes were stained with WGA-488 (second column) and membrane permeability was monitored with PI (third column). proMbxA did not induce membrane damage while MbxA induced formation of spherical membrane protrusions and permeabilization. Growth of spherical membrane protrusions and permeabilization are highlighted by the white boxes in the second and third column and shown in the insets in the right most column as a merge of WGA-488 (green) and PI (red) fluorescence (Scale bar 50 µm, inset 10 µm).

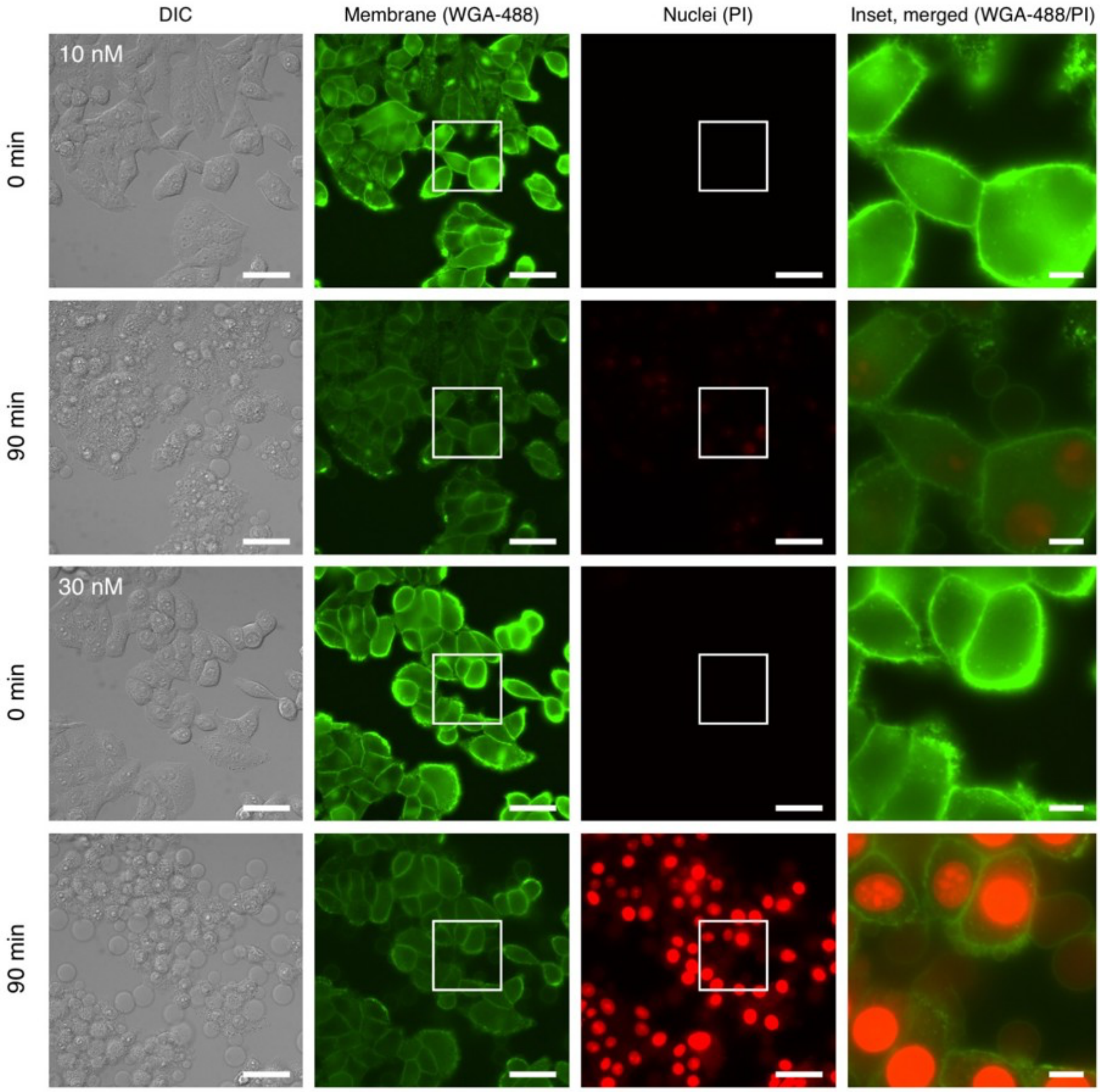


**Fig. S5**. Live-cell imaging of HEp-2 cells treated with 10 nM (first and second row) or 30 nM of MbxA (third and fourth row) over a duration of 90 min. HEp-2 cells of the experiment shown in Fig.S4 after 0 min and 90 min of incubation. Membranes were stained with WGA-488 (second column) and membrane permeability was monitored with PI (third column). Visible staining after extended period of time verifies the potential permeabilizing effect at lower concentrations of MbxA, but with different strength. Growth of spherical membrane protrusions and permeabilization are highlighted by the white boxes and shown in the insets on the right as a merge of WGA-488 (green) and PI fluorescence (red) (Scale bar 10 µm).
